# BAG6 and RNF126 promote the degradation of cytosolic misfolded proteins that contain buried degron motifs

**DOI:** 10.64898/2026.04.24.720685

**Authors:** Sahil Chandhok, Heather A. Baker, Aastha Dalal, Oliver Hong, Aubrey Tierney, Darasimi Kola-Ilesanmi, Vivian Mo, Chase Talarico, Elizabeth Hui, Jonathan P. Bernardini, Shriya Kamat, Calvin Yip, Thibault Mayor

## Abstract

Missense mutations account for the majority of catalogued human disease-associated variants, and many are predicted to destabilize proteins and promote their degradation. To characterize the pathways responsible for recognizing and clearing such variants, we employed a two-pronged approach to identify both quality control components mediating turnover of misfolded proteins and the sequence elements within their substrates that drive this process. Using a panel of unstable cytosolic missense variants in proximity-labeling and RNAi-based experiments, we identified the BAG6-RNF126 pathway as contributing to the clearance of a subset of these substrates. Applying a tile-based approach to a model cytosolic protein, we uncovered strong potential degrons, including a C-terminal degron degraded in part in a BAG6- and RNF126-dependent manner. Modeling supports that this degron can be accommodated by BAG6. Together, our findings add to the growing body of evidence implicating the BAG6-RNF126 pathway as a key mediator of cytosolic protein quality control.

## Introduction

How cells determine whether a misfolded cytosolic protein should be refolded or degraded remains poorly understood. Premature degradation risks eliminating potentially functional proteins, whereas delayed clearance can lead to aggregation and toxicity^1^. This “triage” decision becomes especially critical in disease contexts: during aging, declining chaperone and proteolytic capacity promotes protein aggregation^2–6^, while in cancer, the increased burden of passenger mutations and aneuploidy heightens dependence on protein homeostasis (i.e. proteostasis)^7–9^.

Mutations can induce structural alterations that are detected by the proteostasis network, directing affected proteins toward degradation^10^. This is particularly relevant given that, according to the 2024.4 release of the Human Gene Mutation Database, more than 275,000 missense mutations are associated with human diseases or genetic disorders^11^. Furthermore, a recent deep mutagenesis screen of 500 human protein domains revealed that up to 60% of missense variants are degraded in cells^12^. Despite this, we lack mechanistic insight into how misfolding alters recognition by the folding and degradation machinery. This challenge is compounded by the scale of the human proteostasis network, which includes nearly 200 chaperones and more than 600 E3 ubiquitin ligases^13–15^, many of which could influence triage decisions.

To elucidate the targeting mechanisms involved, we previously characterized single missense variants of several cytosolic proteins associated with genetic diseases and known to undergo proteolysis ^16^. In all but one case, degradation of these mutants in tissue culture cells was dependent on ubiquitination and proteasomal activity. Surprisingly, knockout (KO) or knockdown of the STUB1/CHIP ubiquitin ligase, the first E3 proposed to target misfolded cytosolic proteins for degradation^17,18^, did not affect the turnover of any of the tested variants^19^. Similarly, individual KOs of the HUWE1, UBR4, UBR5, UBE2O or HERC1 E3 ligases, each previously implicated in targeting misfolded or unassembled cytosolic proteins, did not markedly affect the degradation of three representative variants we assessed^19^. These findings point to the existence of additional, yet unidentified, pathways for targeting misfolded cytosolic proteins to the proteasome.

Proteins are targeted for degradation though the recognition of short sequence elements termed degrons^20^. A prevailing model of protein quality control posits that many proteins harbour buried degrons that become exposed upon misfolding, thereby enabling recognition by the ubiquitin proteasome system (UPS)^21^. In yeast, the E3 ligases San1 and Rsp5 have been shown to recognize exposed hydrophobic regions or residues following misfolding and heat stress^22,23^. Similarly pathological mutations in DHFR (dihydrofolate reductase) induce protein misfolding and exposure of degradation promoting elements^10^. More recently, large-scale studies have identified ∼60,000 degrons in the human proteome^24,25^. The fact most of these sequences are normally buried underscores their potential importance in protein quality control. However, it remains unclear whether a dominant pathway exists that both recognize such exposed degrons and mediated the degradation of cytosolic proteins misfolded by missense mutations.

In this study we used proximity labeling–based proteomics to identify components of the proteostasis network that interact with cytosolic proteins carrying missense mutations. Subsequently, using RNAi-based experiments, we identified the BAG6–RNF126 pathway as being involved in the turnover of a subset of the mutant proteins analyzed. Further mapping led to the identification of several degron sequences in one mutant protein, including a C-terminal degron that is degraded in a BAG6- and RNF126-dependent manner. Thus, in this study, we establish the BAG6–RNF126 pathway as an important contributor to the selective degradation of cytosolic proteins destabilized by missense mutations.

## Results

### Unstable cytosolic missense mutants display increased interactivity with members of the proteostasis system

We first sought to expand the BioID proximity-labelling approach to more broadly identify components of the protein homeostasis network that triage cytosolic misfolded proteins for degradation. To this end, we included seven unstable mutant variants derived from six different gene products. These cytosolic proteins, which contain destabilizing missense mutations, are degraded by the UPS^16^.

For the BioID experiments, we C-terminally tagged either the wild-type or unstable mutant variants with the TurboID biotin ligase and stably expressed these constructs in HEK293T Flp-In cells (Figure 1A). As a control, we similarly tagged and expressed *Renilla* luciferase (RLuc). We first confirmed the expression of all the baits following a 24-hour tetracycline induction. As expected, mutant proteins accumulated to lower steady-state levels than their corresponding wild-type counterparts (Figure S1A-C), indicating the addition of the TurboID tag did not stabilize the mutants or interfere with their degradation. To verify that TurboID tagging did not affect cytosolic localization, we performed immunofluorescence microscopy and confirmed that all TurboID-tagged reporters retained their expected cytosolic localization (Figure S2).

**Figure 1.**
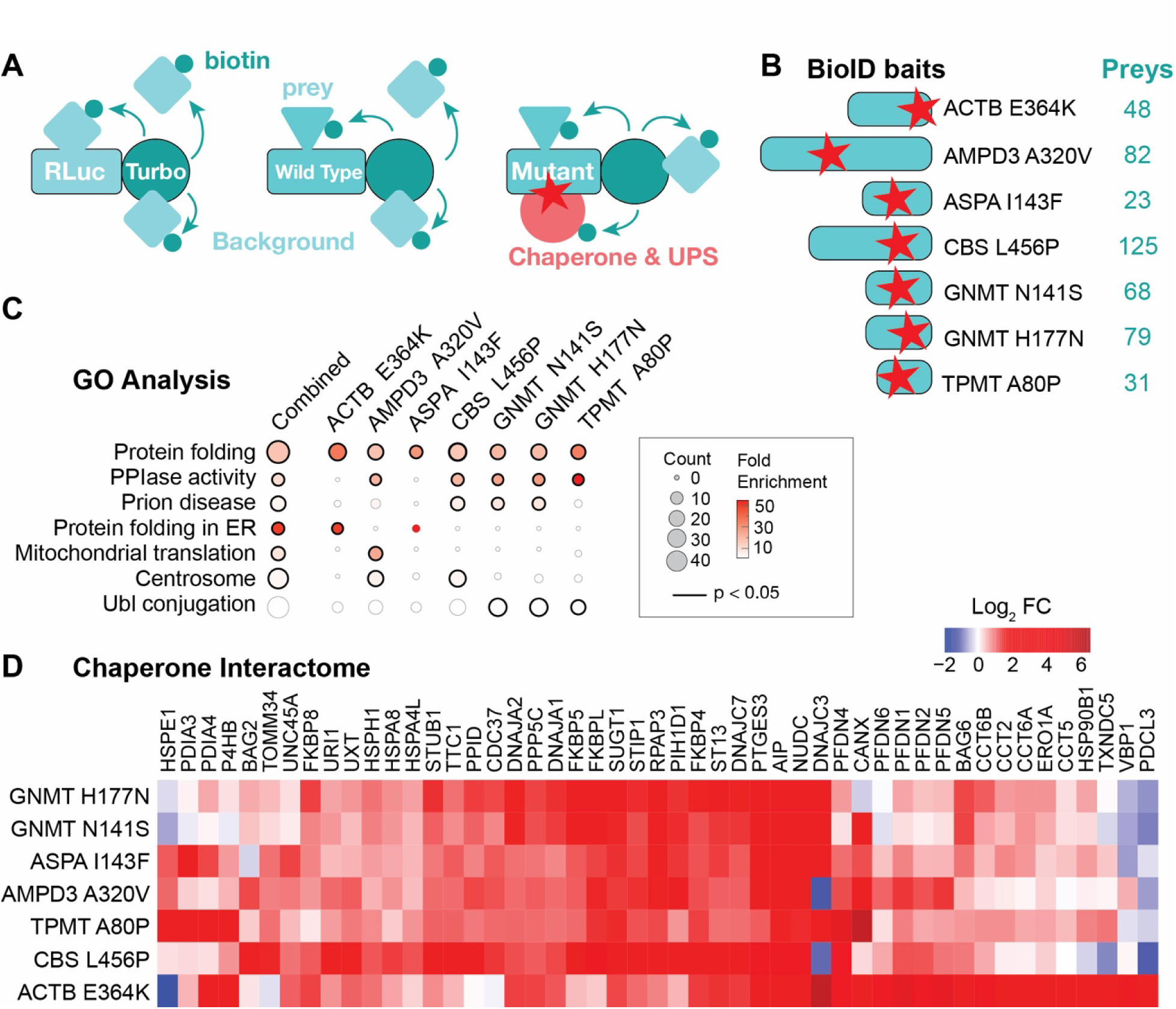
BioID analysis of cytosolic misfolded mutants reveals an extensive network of interacting chaperones. **A.** BioID schematic. **B.** Summary of the number of high-confidence prey for each mutant. **C.** GO analysis of proteins enriched with the indicated baits compared to the corresponding wild-type proteins. Only significant and non-redundant GO were retained (after Benjamini p value corrections). **D.** Heat map of chaperones showing high interaction scores (SAINTq) and enrichment in mutant proteins relative to the corresponding wild-type proteins, defined by a >2-fold change (FC) and *t*-test significance in at least one of the assessed baits (n = 2 technical replicates for 2 biological replicates).

To identify mutant-specific interacting proteins, we carried out a 30-minute biotin labeling followed by cell lysis, automated streptavidin-based pull-down, and data-independent acquisition (DIA) mass spectrometry analysis. Candidate interactors were first determined using SAINTq, and we subsequently applied a t-test to identify proteins significantly enriched with the mutant variants relative to their corresponding wild-type proteins (Supplementary Table 1). Across all the datasets, we identified between 23 and 125 proteins specifically enriched with mutant baits (Figure 1B, S3A). Of these, 54 proteins were enriched in at least three mutants, for a total of 244 proteins enriched in one of the BioID experiments (Figure S3B).

Gene ontology analysis revealed that proteins involved in protein folding were enriched across all mutant datasets (Figure 1C). Notably, 51 distinct chaperone proteins were enriched in at least one mutant, and most of these were also identified in a second independent BioID experiment (Figure S3C-D, Supplementary Table 2). The majority of these chaperones have been reported to localize in the cytosol (Figure S3E). When integrating results across all the baits, a subset of chaperones, such as DNAJA2, displayed increased association with nearly all mutant proteins (Figure 1D). These observations suggest that core elements of the proteostasis network engage broadly with structurally destabilized cytosolic proteins to maintain protein quality control.

In contrast, other chaperones show more selective enrichment with specific mutant proteins. For instance, a mutation in β-actin caused enhanced association with the TRiC chaperonin and prefoldin complexes (Figure 1D), which are known to facilitate actin folding^26^, and is consistent with recent single-molecule tracking analysis^27^. In addition to chaperones, we also identified several components of the UPS, including multiple E3 ligases, that were preferentially enriched with several mutant proteins relative to their wild-type counterparts (Figure 1C, S3F). These results indicate cytosolic misfolded proteins are recognized by a broad spectrum of elements of the proteostasis network, spanning molecular chaperones and the protein degradation machinery.

### RNAi-based knockdown identifies BAG6 and RNF126 as key players in the turnover of cytosolic misfolded proteins

To determine which components of the proteostasis network play dominant roles in targeting cytosolic misfolded proteins for degradation, we screened over 30 candidates by RNAi. These experiments used HEK293T cell lines, each stably expressing one of four mutant substrates via a dual-fluorescence reporter system. In this system, mutant proteins were C-terminally tagged with a green fluorescent mClover2 protein, followed by a P2A self-cleaving peptide and a red fluorescent mRFP1 protein (Figure 2A). Measuring the mClover2-to-mRFP1 fluorescence ratios by flow cytometry provides a readout of mutant protein stability, while normalizing for expression and translation levels. The RNAi experiments broadly indicated that the knockdown of chaperones tended to have a destabilizing effect on mutant reporter levels, highlighting their role in the promotion of client stability (Figure 2B). Interestingly, except for BAG6 RNAi-treated samples, knockdown other UPS components such as the STUB1 E3 ligase had little to no effect on the stability of the reporter substrates, again pointing to potential functional redundancy among quality control degradation pathways.

**Figure 2.**
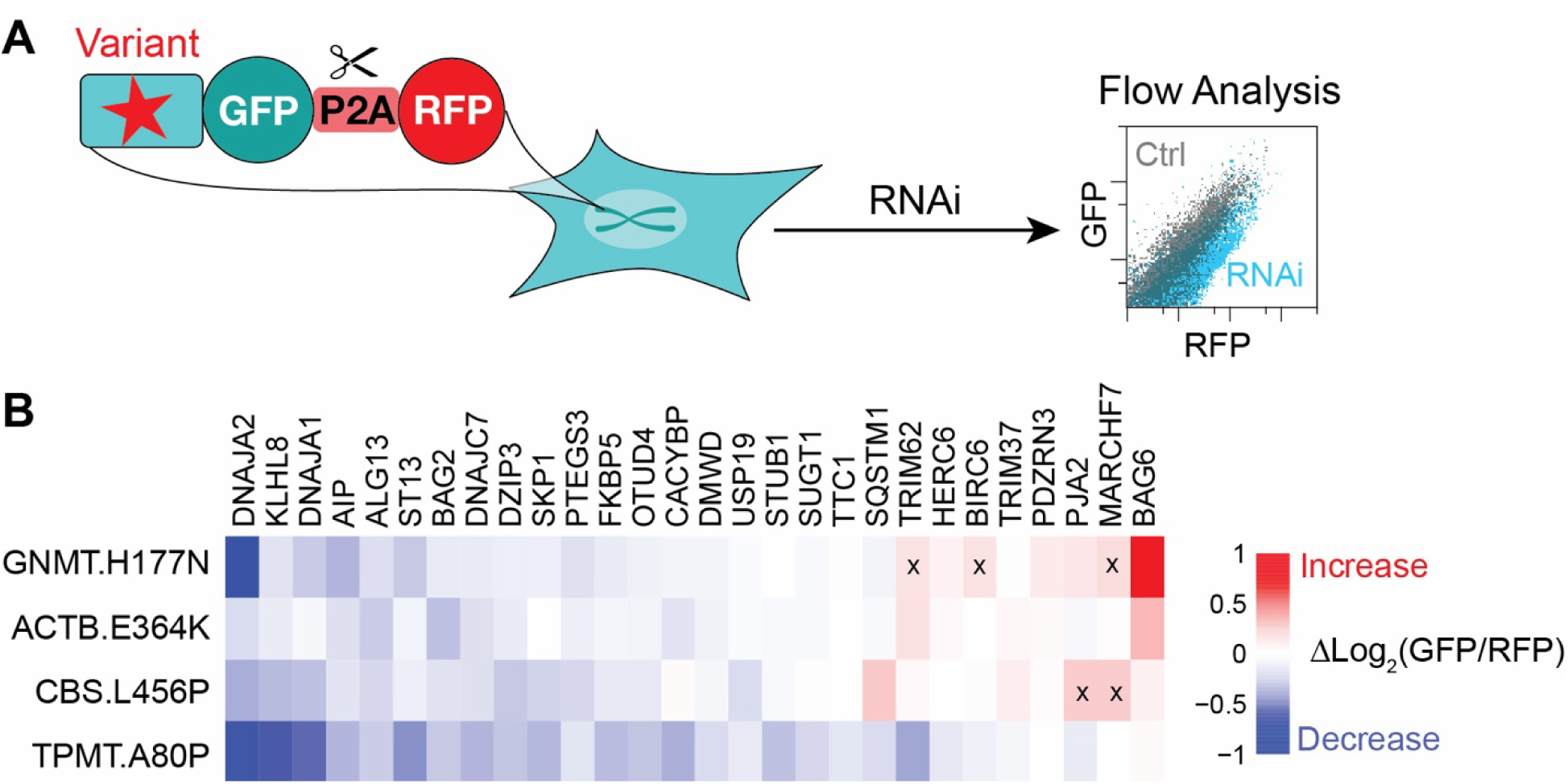
Functional screening of BioID hits by RNAi. **A.** Schematic of the P2A construct transduced into HEK293 cells for the RNAi experiments. Four different variants were C-terminally tagged with mClover2 (GFP). RFP was used to normalize the signal following flow cytometry analysis. **B.** Heat map of flow cytometry analysis of the indicated reporters transfected with 31 different RNAi constructs. The RFP-normalized GFP signal was compared to control RNAi for each mutant (*n* = 1). “×” denotes samples in which RFP levels also appeared to be affected by the RNAi.

BAG6 is a cytosolic and nuclear chaperone of particular interest to us, as it displayed a marked increase in association with the β-actin variant and two glycine N-methyltransferase (GNMT) variants, concomitant with the stabilization of these mutant proteins upon knockdown. Conserved in metazoans, BAG6 contains a N-terminal ubiquitin-like (UBL) and BUILD domains, a C-terminal BAG domain, and two less-characterized central regions (domain II and proline-rich region), which are predicted by AlphaFold to form distinct domains (Figure 3A). BAG6 was initially shown to facilitate the targeting of tail-anchored (TA) proteins via its BAG domain and interactions with members of the GET (Guided Entry of TA pathway) complex^28–31^. In contrast, BAG6 also mediates proteasomal degradation of mislocalized transmembrane proteins through association with the RING finger E3 ligase RNF126^32^. Subsequent studies implicated BAG6 in the degradation of aberrant translation products and the aggregation-prone TDP-43 protein associated with amyotrophic lateral sclerosis^33–35^.

**Figure 3.**
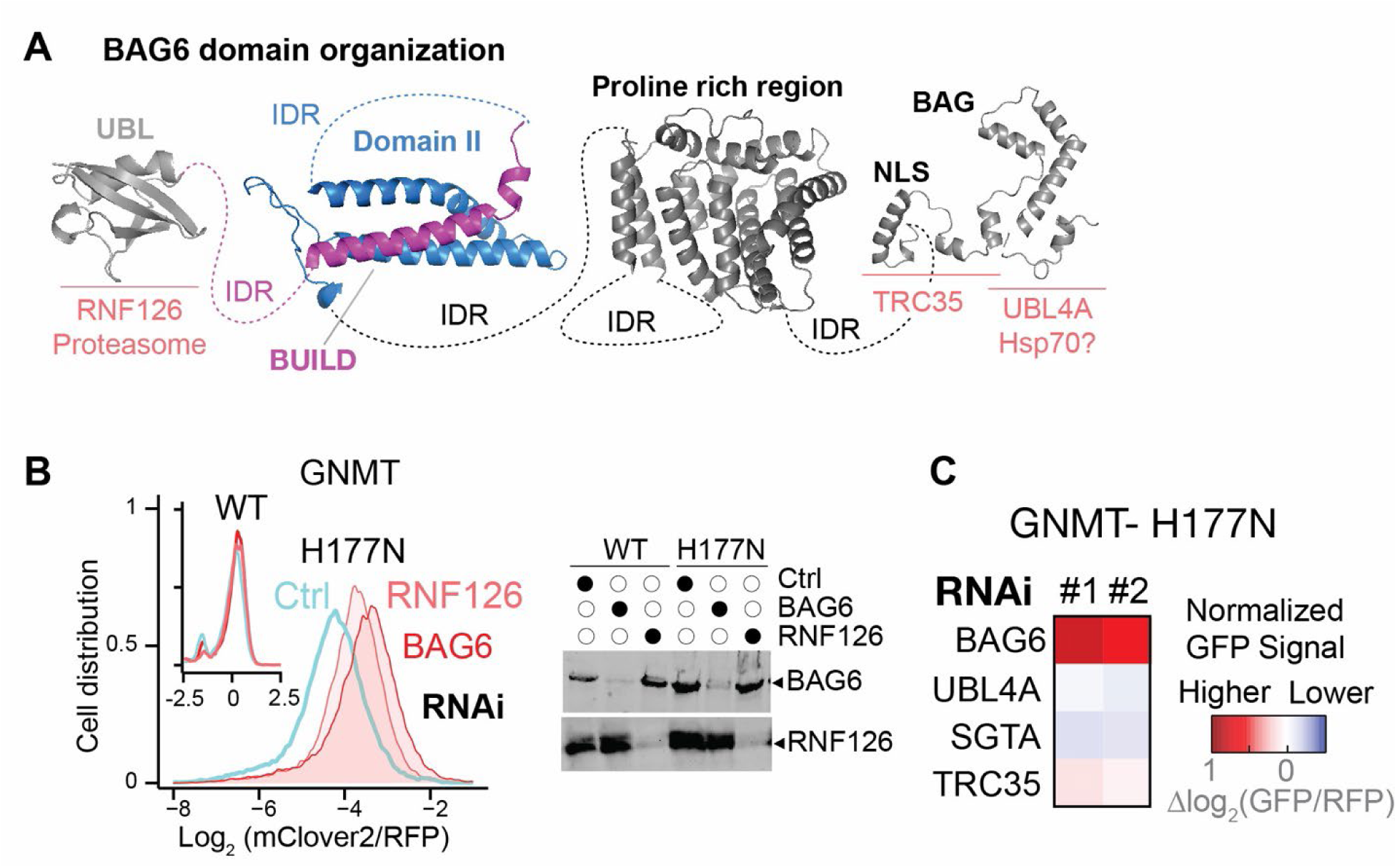
BAG6 and RNF126 promote degradation of the mutant cytosolic GNMT protein. **A.** Schematic of Bag6 domain topology (IDRs not to scale) and proteins known to bind each region. The C-terminal portion of BUILD and Domain II are predicted to form a structured domain, and residues at the end of the magenta α-helix have been shown to be important for binding hydrophobic elements^44^. **B.** Flow cytometry analysis of cells transduced with WT or mutant GNMT reporters and transfected with the indicated RNAi, along corresponding Western blots (n=1). Two additional independent experiments are shown in Figure S4A and loading control in S4B. **C.** Averaged normalized GFP signal of cells expressing the transduced mutant GNMT reporter transfected with the indicated RNAi and compared to control RNAi (n=3 for each 2 oligos from a different provider than in panel B; detailed data shown in Figure S4C). * indicates knockdowns that could not be validated by mass spectrometry.

To validate the results of our RNAi screen, we used siRNA from another manufacturer and independently confirmed the role of BAG6 in the turnover of the GNMT-H177N mutant protein, while additionally showing that wild-type GNMT levels are not impacted (Figure 3B, S4A, B). Furthermore, we found that RNF126 also played a specific role in the degradation of the misfolded GNMT variant (Figure 3B, S4A). In contrast, knockdown of other components of the GET complex (UBL4A, TRC35, SGTA) had no effect on either mutant or wildtype stability (Figure 3C, S4C). Together, these data support a role of BAG6 in the turnover of cytosolic misfolded proteins and indicate that this process is mediated through the RNF126 E3 ligase.

### Tile-based scanning uncovers functional UPS-targeted degrons within GNMT

Recent work has identified short sequence motifs within proteins that can act as degrons^25^ and it has been proposed that cytosolic misfolded proteins are targeted for degradation upon exposure of normally buried degrons^36^ Therefore, we next focused on identifying such sequences within the full-length GNMT protein. To achieve this, we generated 28 partially overlapping 20-mer peptide tiles (with a 10 amino acid overlap), one 15-mer tile at the C-terminus, and three additional tiles encompassing disease-associated substitutions (L50P, N141S, H177N) (Figure 4A). Because many degradation signals are short linear elements^20^, this approach focuses on sequence-encoded signals rather than predicted secondary structure. Peptides were fused to bicistronic fluorescent reporters to assess the stability of each GNMT fragment (Figure 4A). The fluorescent construct used here consists of DsRed followed by an internal ribosomal entry site (IRES) and EGFP fused to the N-terminus of the tile sequence. Since both fluorescent proteins are on the same mRNA, DsRed serves as an internal control for expression, while EGFP provides a proxy for stability of the fusion protein. An additional N-terminal tail comprised of glutamate and phenylalanine was also included to prevent the formation of C-terminal degrons^37,38^.We first assessed normalized EGFP levels following transfection of each tile in HEK293T cells by flow cytometry. As a control, we expressed EGFP without any C-terminal sequence. Across the library, we observed a spectrum of levels ranging from no degradation to nearly complete loss of signal (<5% EGFP/DsRed signal remaining) relative to the control (Figure 4B). Notably, five tiles led to strong degradation (≤20% signal remaining) and six caused moderate degradation (≤50% signal remaining) out of 32 tested. Strong degron-containing tiles were enriched within the N- and C-terminal regions (within the first and last 60 residues) and partially overlapped with both α-helices and β-strands in the native structure (Figure S5A).

**Figure 4.**
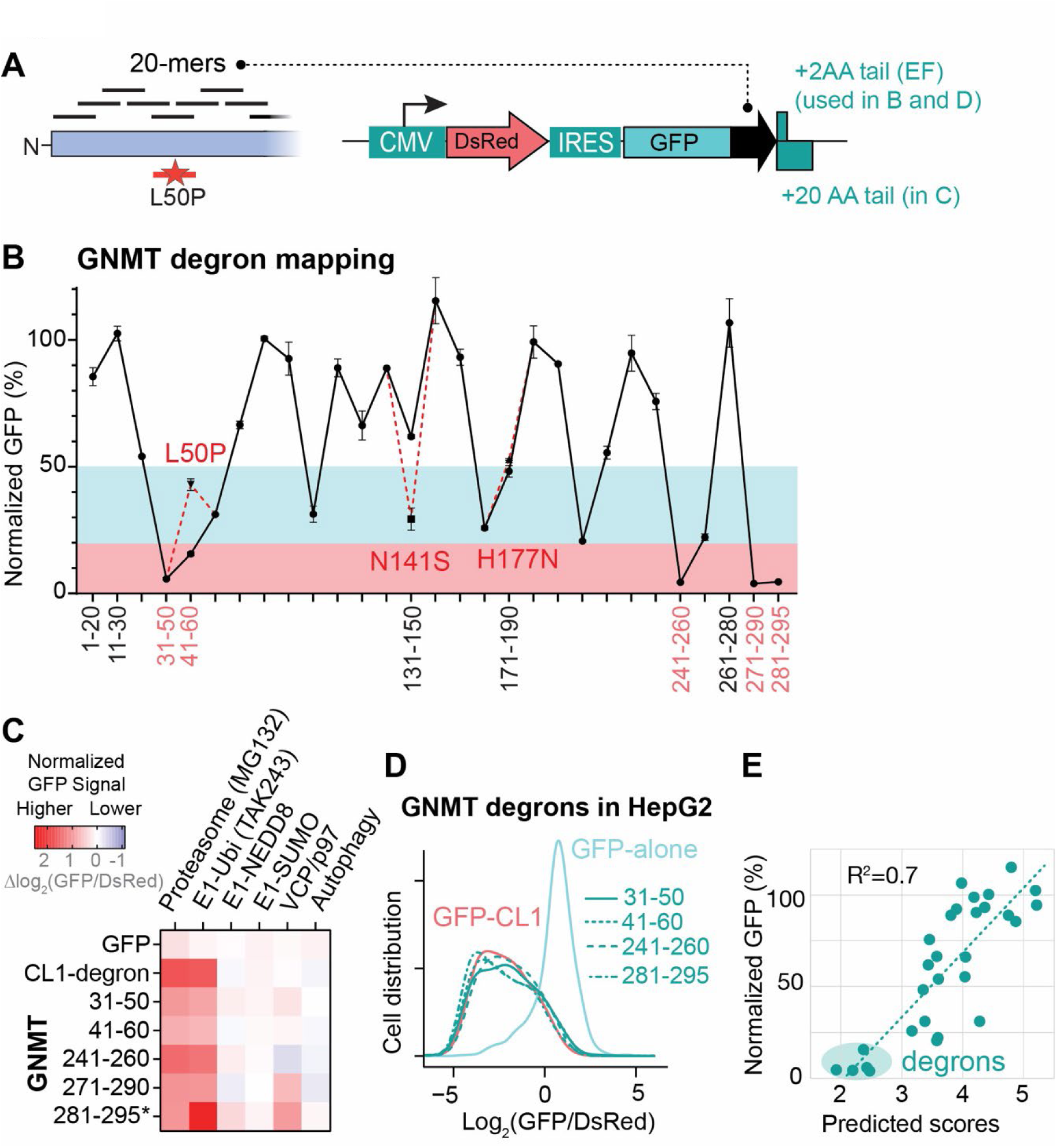
GNMT contains distinct regions recognized by the UPS. **A.** Schematic of the tiling approach using partially overlapping GNMT 20-mers fused to GFP in an IRES plasmid, together with constant C-terminal residues to prevent the formation of artificial C-end degrons. **B.** GFP signal normalized to RFP following transient transfection in HEK293 cells and flow cytometry analysis of the indicated GNMT fragments (n=3). **C.** Normalized GFP signal in HEK293 cells transduced with the indicated GNMT degrons after 16 h treatment with inhibitors targeting the indicated enzymes or cellular processes (n=2 or 3; data expanded in Figure S5C). CL1 serves as a positive degron control, and * denotes a STOP codon instead of an added C-terminal tail. **D.** Expression of GNTM degrons in HEPG2 cells. Representative histogram of the indicated plasmids containing GNMT degrons or controls were transfected into HepG2 cells prior to flow cytometry analysis (n=3, data expanded in Figure S5D). **E.** Comparison between GNMT degron mapping results shown in panel B and predicted degron scores.

Disease-associated mutations of GNMT had varied effects on peptide stability. Outside of the context of the full-length protein, only the N141S disease-associated substitution resulted in a marked increase in tile degradation. In contrast, the L50P mutation in degron 41-60 reduced degradation relative to the wild-type sequence, whereas the H177N had no effect on tile 171-190 (Figure 4B). These findings are consistent with recent work showing that only a subset of mutations generate degron motifs^24^. Overall, these observations suggest that mutation-induced degradation is likely initiated by protein misfolding, which subsequently exposes normally buried degrons that can be recognized by cellular quality control machinery.

To assess context-dependent accessibility, we retested the strong and moderate degrons using a second reporter that includes a 20-amino acid C-terminal extension (Figure 4A). This extension allows us to test whether each degron remains functional when positioned internally within a longer polypeptide chain, as would occur in a misfolded or partially folded protein. The additional C-terminal sequence thus mimics the structural context of degrons embedded within full-length cytosolic proteins. These constructs were transduced in HEK293 cells for stable expression. Although most degrons showed some stabilization with the added C-terminal tail, the overall ranking of degron strength was largely preserved, and all strong degrons maintained robust degradation (Figure S5B). Notably, the 281-295* tile, which retains the natural stop codon, was assessed without any C-terminal sequence to preserve its native context and displayed similarly strong degradation. These results indicate that these sequences function as internal degrons rather than artefacts arising from C-terminal exposure.

We next sought to determine whether degradation mediated by the strong GNMT degrons depends on the UPS, as observed for full length GNMT mutants^16^. We tested a panel of small-molecule inhibitors targeting major degradation pathways. As a positive control, we included the 16-residue CL1 degron, which has been shown to be degraded by the UPS^39^.The proteasome inhibitor MG132 and the E1-activating enzyme inhibitor TAK-243 stabilized all strong degrons and CL1, consistent with UPS-mediated degradation (Figure 4C, S5C). Degrons 281-295 and 281-295* were modestly stabilized by VCP/p97 inhibition with CB-5083. Blocking either Hsp70 or Hsp90 also marginally stabilized the 281-295* degron. In contrast, the autophagy inhibitor Bafilomycin A1 did not appear to have any significant effect on degron stability. Moreover, blocking the NEDD8-activating enzyme with MLN4924 had no impact, suggesting that turnover of these degrons is independent of Cullin-RING ligases (CRLs) that require neddylation for activity^40^. Collectively, these results indicate that degradation of the identified GNMT degrons is mediated predominantly by the UPS.

GNMT is highly expressed in liver cells and makes up 1-3% of total liver proteins across different species (REF). To confirm that the strong GNMT degrons are also recognized in physiologically relevant cellular context, we tested them in HEPG2 hepatocellular carcinoma cells. Consistent with GNMT’s hepatic expression profile, the degrons remained destabilized in HEPG2 cells (Figure 4D, S5D), indicating that the QC pathways targeting GNMT degrons appear to be preserved across cell types.

Remarkably, degradation-prone regions within a linear cytosolic protein sequence can now be predicted with good accuracy^24,25,41^. For example, a predictor developed by the Elledge lab^25^, which we refined using additional parameters (Figure S5E), shows a strong correlation between predicted degron scores and the measured levels for each GNMT fragment (Figure 4E). These results demonstrate that GNMT contain several discrete sequence elements capable of driving degradation via the UPS.

### Residue-level mapping defines GNMT degrons and implicates BAG6–RNF126 in their recognition

To assess the presence of motifs contained within GNMT degrons, we performed scanning mutagenesis across five segments. Each position was substituted with amino acids of differing properties, similar to Zhang et al. (2023). In this scheme, bulky hydrophobic residues (F, M, P, W) were replaced with serine; small or aliphatic residues (A, C, D, E, G, I, L, S, T, V) were replaced with arginine; and polar or charged residues (H, K, N, Q, Y) were replaced with alanine. Using this approach, we pinpointed stabilizing substitutions across each of the assessed tiles to identify specific residues seemingly important for degron activity (Figure 5A).

**Figure 5.**
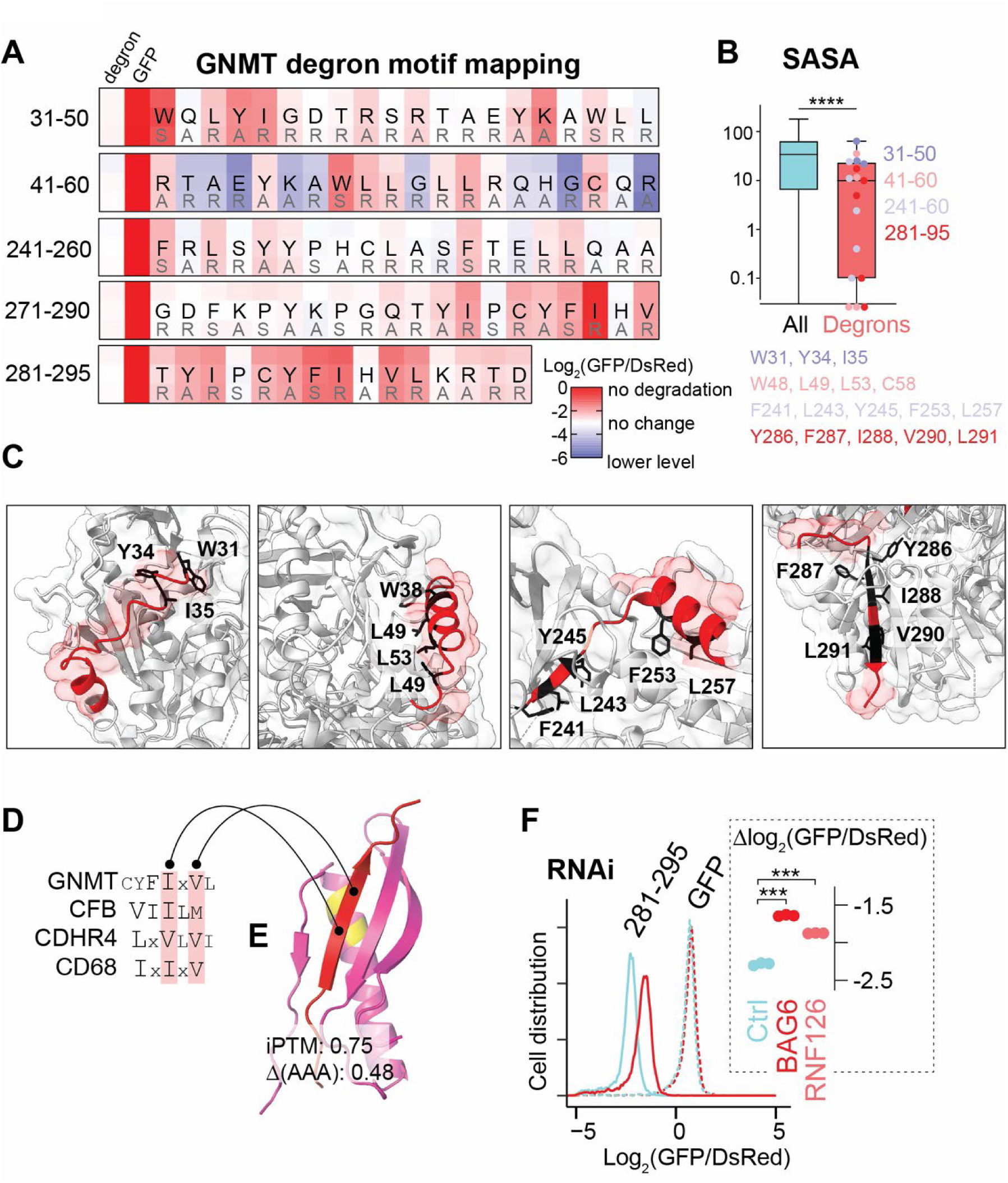
GNMT contains distinct normally buried degrons, including a C-terminal sequence degraded in a BAG6- and RNF126-dependent manner. **A.** Normalized GFP signal following mutation of the indicated GNMT degron residues (to A, R, or S) on 5 tile regions after transient transfection and flow cytometry analysis (n=3; each mutation position was assessed individually). **B.** Box plots of the solvent-accessible surface area (SASA) of GNMT residues in the tetramer (1R74; square Angstroms). An unpaired t test with Welch’s correction was done to compare all residues with key degron residues (p value < 0.0001). **C.** Mapping of the indicated key degron residues onto the GNMT tetramer (1R74). **D.** The putative BAG6 degron motif located within β-strand sequences. **E.** AlphaFold3^58,59,62,63^ prediction of the GNMT degron (red) docking onto the BAG6 BUILD domain (hydrophobic residues shown in yellow) and forming a β-strand within the IDR (not observed with the inactive AAA degron). **F.** HEK293 cells expressing a degron-tagged GFP (GNMT residues 281–295) and DsRed were subjected to RNAi-mediated knockdown of BAG6 (red) or RNF126 (pink), with a non-targeting scramble control (Ctrl, blue). Representative histogram of normalized GFP/DsRed signal following the indicated RNAi treatments. Inset shows quantification of Δlog₂(GFP/DsRed) relative to GFP control. Mean values with s.d. are shown; statistical significance was determined by unpaired t-test (n = 3; *** p ≤ 0.001, **** p ≤ 0.0001; data expanded in Figure S6D).

Although the GNMT 31–50 and 41–60 regions contain overlapping sequences, they each harbor distinct clusters of key residues that exert strong stabilizing effects when mutated, thereby defining two separate degrons. In GNMT 31–50, residues W31, Y34, I35, and K46 had the greatest impact on degron activity (Figure 5A). However, mutation of the first three residues did not fully block degradation of the tile, consistent with the presence of a second degron within this region (Figure S6A). In GNMT 41–60, W48, L49, L53, and C58 were the strongest stabilizing residues, and mutation of the first three was sufficient to nearly abrogate degradation (Figure S6A). In contrast, multiple residues distributed across the GNMT 241–260 region each conferred moderate stabilization when mutated, suggesting a longer degron with two clusters of key residues. Mutation of four residues (F241, L243, F253, and L257) was required to fully inactivate this degron (Figure S6A). The C-terminal degrons displayed a stretch of hydrophobic residues that strongly stabilized the reporters when mutated. Specifically, mutation of F287, I288, V290, and L291 markedly stabilized both GNMT 271–290 and 281–295 segments (Figure 5A). Because the same residues influence degradation in both tiles, these regions likely contain the same degron motif, and mutation of the first three residues nearly abolished degradation (Figure S6A). Collectively, mapping across these tiles identifies four strong degrons within the GNMT sequence.

All the residues identified above are buried in the native GNMT structure and less solvent accessible (Figure 5B, C), except for K46. The contribution of K46 may reflect its role as a ubiquitination site in the context of the assessed tile. This is consistent with the proposed tripartite degron model, which involves a degron sequence and a nearby ubiquitinated lysine^42^. While not all strong degron tiles contain a lysine, EGFP contains multiple lysine residues that could serve as ubiquitination sites. Consistent with the importance of these degron residues, their mutation in the context of full-length GNMT led to stabilization of the protein, with the exception of the first degron that is also potentially more accessible in the native structure (Figure 5B, S6B, C). Together, these results indicate that GNMT contains several potent degrons composed of residues that are normally buried in the native structure and could become exposed upon misfolding.

Several recent studies have systematically catalogued degron motifs and their cognate E3 ligases, including large-scale stability reporter screens and curated databases such as DegronID and DEGRONPEDIA^25,43^. We used these resources to compare the GNMT degron sequences with known motifs and infer potential recognition factors. The GNMT 31–50 degron resembles the W[VAC]x[YRT][ILT] motif recognized by FEM1B, an adaptor of the CUL2 complex^25^; however, FEM1B RNAi did not significantly stabilize this degron (Figure S6D). Mutation of this degron in the full-length variants also did not stabilize the mutants (Figure S6C), indicating that this degron alone cannot account for the degradation of the misfolded proteins. The GNMT 241–260 region shares features with the amphipathic helix motif (fxxffxxfxxxfxxf) proposed to be recognized by FBXO21^25^. However, structural mapping shows that the key residues span both a β-strand (241–245) and an α-helix (249–260), suggesting a different mode of recognition. Consistently, FBXO21 RNAi had no effect on degron stability (Figure S6D). Together with the lack of stabilization upon MLN4924 treatment (Figure 4C), these results argue against a major role for Cullin-RING ligases in GNMT degron turnover.

Two GNMT degrons show features consistent with BAG6 recognition. The GNMT 45–53 region lies within an α-helix and contains a repeated leucine pattern reminiscent of previously described BAG6-binding motifs^25^. In addition, three other BAG6-dependent degrons recently identified^25^ share sequence similarity with the GNMT C-terminal degron (Figure 5D), and in each case these residues reside within β-strands in the native structure. We propose that the C-terminal degron may engage BAG6 through β-sheet pairing, stabilized by backbone hydrogen bonding and hydrophobic interactions. Consistent with this model, AlphaFold3 predicts that the GNMT C-terminal degron, but not an inactive mutant, docks against hydrophobic residues within the C-terminal region of the BUILD domain (Figure 5E), previously implicated in substrate recognition^44^. In agreement with a potential central role of the GNMT C-terminal degron, depletion of BAG6 or RNF126 impaired its degradation (Figure 5F, S6E). In addition, BAG6 knockdown also significantly—albeit modestly—stabilized the other three degrons, whereas RNF126 depletion had little to no effect on these regions (Figure S6D). This suggests that BAG6 may act more broadly in degron recognition, potentially by facilitating delivery of hydrophobic substrates to the proteasome through its UBL-mediated interactions with the 19S regulatory particle^45,46^. Importantly, our RNAi analyses consistently identify BAG6 as the dominant factor influencing degron-mediated turnover, with RNF126 contributing more selectively, particularly for the C-terminal degron. Together, these results highlight the BAG6–RNF126 axis as a major pathway for the recognition and degradation of misfolded GNMT, with the C-terminal degron emerging as a key signal driving this process.

## Discussion

Factors involved in the turnover of cytosolic misfolded proteins remain to be fully characterized. Here, we have utilized a series of cytosolic mutant misfolded proteins in proximity labelling experiments to uncover nearly seventy different quality control elements that display increased interactivity with at least one of the mutant proteins. Subsequent RNAi-based experiments showed that knockdown of BAG6 and RNF126 impairs the degradation of mutant GNMT but has no impact of steady-state levels of the wild-type protein. Applying a tile-based screening approach to GNMT led to the identification of four effective degrons. Further RNAi experiments indicated that the C-terminal degron is also, at least partially, degraded in a BAG6- and RNF126-dependent manner.

Though the mutant substrates display increased interaction with numerous chaperones, RNAi-mediated depletion of several of these elements resulted in little to no impact on mutant stability, highlighting the potential functional redundancy in these quality control pathways. This was true for instance of KD of the FKBP5 peptidylprolyl isomerase and Tetratricopeptide repeat protein 1 (TTC1), although we could not individually assess each RNAi efficacy in this screen. Nonetheless, several depleted chaperones led to further destabilizing phenotype for the mutant proteins. This observation is in line with the results of our recent study^16^, wherein we have showed that DNAJA2 and A1 could play a role in buffering the degradation of some of the assessed misfolded variants.

Similarly, while we identified several E3 ligases and deubiquitinating enzymes (DUBs), their knockdown had no or only minor impact on reporter levels, potentially due to pathway redundancy. Future studies should therefore consider combinatorial approaches in which multiple candidates are targeted simultaneously. For example, depletion of several key DUBs antagonizing quality control E3s could accelerate degradation of misfolded proteins, consistent with previous work^47,48^. Intriguingly, RNF126 was not identified in the BioID experiment, which may be due to its low lysine content, potentially limiting its biotinylation efficiency.

Using our tile-based approach, we identified four motifs that could function as degrons within the full-length GNMT protein, and these sequences were concentrated in the N- and C-terminal regions of the protein. Furthermore, we also found that destabilizing pathological mutations had little to no effect on tile stability, indicating that the destabilizing consequences of these mutations likely manifests through misfolding and exposure of degradation promoting elements. Indeed, all key degron residues are buried in the native structure, with the exception of the first degron, which is located in a flexible region involved in substrate entry^49^.

Similar to full-length GNMT, the C-terminal degron was degraded in a BAG6- and RNF126-dependent fashion. Based on AlphaFold modeling, the C-terminal degron may bind to disordered region of the BUILD domain of BAG6 forming additional β-strands. One possibility is that this disorder-to-order transition positions the GNMT substrate closer to the UBL domain of BAG6, which recruits the RNF126 E3 ligase and thereby promotes ubiquitination. This interaction may be reinforced by additional, low affinity contacts with weaker degrons, increasing overall avidity. In contrast, binding of helical TA domains may be mediated by a different region of BAG6 and, in the absence of additional contacts, may instead promote the transfer of the TA protein to other components of the GET pathway.

Consistent with our findings, Pedersen and colleagues have shown that BAG6 and RNF126 contribute to the turnover of cytosolic misfolded proteins, first using an extensive library of missense parkin variants, and then by assessing the turnover of FLCN and PAH missense mutants^50^. In all cases, the degradation-promoting properties of BAG6 and RNF126 were specific to the mutant proteins and not observed for the wildtype counterparts. Together with prior work from Zhang and colleagues showing a broad role for BAG6 in degron recognition^25^, these studies strongly implicate the BAG6-RNF126 pathway as a major component of the proteostasis network responsible for clearing cytosolic misfolded proteins.

## Limitations

Our experiments rely on an overexpression-based model, which can lead to mislocalization of a subset of target protein population. For example, several ER chaperones or mitochondrial ribosome proteins were enriched upon expression of ACTB and AMP deaminase 3 (AMPD3), respectively. Future studies will be needed to assess the turnover of these misfolded proteins in their native physiological context. While our model proposes that unfolding events drive degron exposure, additional structural work will be required to confirm this mechanism, particularly to confirm BAG6-degron docking.

## Resource availability

### Lead contact

Further information and requests for resources and reagents should be directed to and will be fulfilled by the lead contact, Thibault Mayor.

## Materials availability

All unique plasmids generated in this study are available from the lead contact. Stable cell lines will be shared based on availability.

## Data and Code availability

- All Raw proteomic data are deposited in the MassIVE repository (First turboID experiment - ftp://MSV000101561@massive-ftp.ucsd.edu; second turboID experiment - ftp://MSV000101563@massive-ftp.ucsd.edu,; MS validation for BAG6 complex knockdown - ftp://MSV000101565@massive-ftp.ucsd.edu)
- All original code is publicly available and has been deposited at https://github.com/SahilCh95/BAG6-RNF126-cytosolic-degradation and https://github.com/Alpine2048/Degron-prediction-by-composition-seq-motifs.
- Any additional information required to reanalyze the data reported in this study is available from the Lead contact upon request.

## Supporting information

Supplemental Table 2 - Data for Second BioID Experiment

Supplemental Table 1 - Data for First BioID Experiment

## Acknowledgments

We thank Drs. Alice Ting, Anne-Claude Gingras and Stephen Elledge for sharing plasmids, Drs. Hyungwon Choi and Ji-Young Youn and Owen Tsai for help with SAINTq analysis, Drs. Mikko Taipale and Rasmus Hartmann-Petersen for discussion, the UBC Flow and Proteomics facilities (especially Jason Rogalski for assistance with mass spectrometry analysis), and members of the Mayor lab for discussions. This work was supported by a grant from the Canadian Institutes of Health Research (CIHR; PJT 495759), S.C. is recipient of a UBC four-years scholarship and H.B. was supported by a CIHR Doctoral Canadian Graduate Scholarship.

## Author contributions

Conceptualization: S.C., H.A.B., A.D., T.M.; Methodology: S.C., H.A.B., A.D., O.H., S.K., C.T., T.M.; Investigation: S.C., H.A.B., A.D., O.H., A.T. D.K.-I., V.M., C.T..; Technical support for experimental work: E.H., J.P.B.; Formal analyses: S.C., H.A.B., O.H., C.T., T.M; Resources: C.Y., T.M.; Visualization: S.C., H.A.B, T.M.; Writing & editing: S.C., H.A.B, T.M..

## Declaration of generative AI and AI-assisted technologies

During the preparation of this manuscript, we used ChatGPT to correct several sections of the text for grammar and flow and we take full responsibility for the content of the publication.

## Star Methods

### Experimental model and study participant details

#### Cell culture

HEK293T (ATCC), Flp-In™ T-REx™ 293 (Invitrogen) and HepG2 (ATCC) cell lines were maintained in Dulbecco’s Modified Eagle Medium (DMEM), high glucose (Gibco) supplemented with 10% fetal bovine serum (FBS) (Gibco) and 1% penicillin-streptomycin (Gibco) at 37°C and 5% CO2. All routine passaging was performed every 2-3 days and carried out with 0.25% Trypsin-EDTA (Gibco). Parental cells and derived cell lines were routinely tested for mycoplasma contamination.

### Method details

#### Plasmid Cloning

The V5-turboID fragment (a kind gift from Dr. Alice Ting; Addgene plasmid #107169)^51^, was subcloned using Gibson Assembly (New England Biolabs) into pCDNA5 destination vectors C-terminally to GNMT-WT and H177N constructs. All other reporters^16^ were subsequently subcloned into the pCDNA5 GNMT-V5-turboID plasmid using Gibson Assembly. All constructs were confirmed using whole plasmid sequencing and listed in Table S3.

The MSCV-CMV-DsRed-IRES-EGFP-DEST vector was a kind gift of Dr. Elledge (Addgene #41941)^52^ and modified by Gibson Assembly to introduce the XhoI/EcoRI restriction sites instead of the DEST region. GNMT tile sequences were synthesized as complementary DNA oligomers (Invitrogen) with XhoI/EcoRI overhangs, annealed, phosphorylated, and ligated into the digested backbone. The GNMT tiles were similarly cloned in the GPS6.0 lentivirus vector^25^, in which the DEST region was replaced with MluI/XhoI sites along the C-terminal extension 20AA (QGRARPNQEVQIGEMENQLS)^25^. To generate the multi-alanine mutations in GNMT, site-directed mutagenesis was used on the full length GNMT and mutant reporter constructs.

#### Cell line generation

Flp-In™ T-REx™ 293 cells were grown to 70% confluence in 35 mm dishes and co-transfected with 2 µg of pOG44 flp recombinase vector (Table S3) and 1 µg of bait-V5-TurboID constructs using FuGENE HD (Promega). The media was replaced the next day, and cells were expanded to a 100 mm dish 24 hours later. Polyclonal cell populations were selected for 15-20 days using 200 µg/ml of Hygromycin B, and the dishes were split in half, into ‘population A’ and ‘population B’(biological replicates). Cells were grown to 70% confluence before 1 µg/ml tetracycline induction for 24 hours in with fresh media. 50 µM biotin (Sigma Aldrich) was added to each dish for either 30 minutes and cells were washed with cold 1x PBS, trypsinized and collected. The cell pellets were lysed in modified RIPA buffer (50 mM Tris-HCL pH=7.4, 150mM NaCl, 0.5% sodium deoxycholate, 0.1% SDS, 1% Triton X-100) supplemented with 1 mM PMSF and 1X Roche EDTA-free cOmplete protease inhibitor cocktail for Western blots and for BioID experiments. Alternatively, cells were grown on poly-D-lysine treated coverslips for methanol fixation and imaging.

To generate stable degron reporters, HEK293T packaging cells were seeded into a 6-well plate, grown in an antibiotic free complete media (high glucose DMEM, 10% FBS), and co-transfected with pMD2.G (0.5 µg; Addgene #12259, http://n2t.net/addgene:12259; RRID:Addgene_12259), psPAX2 (0.5 µg; Addgene #12260), and the viral plasmid of interest (1 µg) using FuGENE 6 (Promega). 24 hours after transfection, the media was discarded and replaced with DMEM, 10% FBS, 1% penicillin/streptomycin. The media was collected and passed through a 0.45 mm filter. HEK293T target cells were seeded into a 6- well plate at a density of 50% confluency and grown in viral media supplemented with 4 μg/ml of polybrene (EMD-Millipore). Transduced cells were selected with 6 μg/ml of blasticidin (Invitrogen) for seven days.

#### Immunostaining and fluorescence microscopy

Flp-In™ T-REx™ 293 cells stably expressing the turboID constructs were seeded on poly-D-lysine treated coverslips. After a 24-hour induction (1 µg/mL tetracycline), the cells were then treated with 50 µM biotin for 30 minutes, fixed for 5 minutes with ice cold methanol, then blocked overnight with 3% bovine serum albumin (BSA, Thermo Fisher Scientific) in PBS. The samples were probed with anti-biotin (Abcam) and V5-tag (Invitrogen) antibodies (1:100 dilution in 3% BSA) for 1 hour at room temperature, washed thrice with 3% BSA, and then probed with fluorescently labelled secondary antibodies (Invitrogen, anti-mouse conjugated to Alexa Fluor^TM^ 594 and anti-rabbit conjugated to Alexa 488 Fluor^TM^, 1:400 dilution) at for 1 hour at room temperature. The samples were then washed thrice with 3% BSA, stained with Hoescht 33342 (Invitrogen, 1:10,000 in PBS) and then mounted on slides with ProLong™ Gold Antifade Mountant (Invitrogen) and sealed. The samples were imaged on an AXIO Observer 7 inverted led fluorescence microscope (Zeiss). Images were captured as a z-stack (34 slices, 3.96 µm) and deconvoluted with the point spread function method. Images were processed and exported as .tif files with the Zen Blue 3.0 image processing software.

#### Sample processing for turboID-mass spectrometry experiments

Cells stably expressing the turboID fusion proteins were seeded in 100 mm tissue culture plates. Once they reached 70% confluency, expression and biotinylation was induced with a 24-hour 1 µg/mL tetracycline and 30-minute biotin treatment, respectively. Cells were washed with cool PBS and scraped into PBS and pelleted (3 min at 1000*g*, 4°C). The supernatant was removed, and cells were snap frozen in liquid nitrogen and stored at -70°C. For cell lysis, pellets were thawed on ice and resuspended in 1 mL of modified RIPA buffer (50 mM Tris-HCl pH 7.4, 150 mM NaCl, 0.1% SDS, 0.5% sodium deoxycholate, 1% triton-X, 1X Roche cOmplete, EDTA-free protease inhibitor cocktail, 1 mM PMSF). The samples were left on ice for 30 minutes, following which they were sonicated in an ice water bath on the highest setting, with a 30 seconds ON/OFF pulse. The lysates were then clarified with a 20-minute centrifugation (16,000 *g*, 4°C). Protein concentration was then assessed with a BCA assay (Pierce) and lysates were snap frozen and stored at -70°C. 2 biological replicates of each cell line were collected (populations A and B).

For the pulldown, lysates were normalized to a concentration of 2.1 µg/µL and for the first bioID experiment, 1 pulldown was performed per biological replicate. The only exception was the *Renilla* luciferase negative control, where 4 pulldowns were performed for the first biological replicate and 2 for the second one. For the second BioID experiment, to increase stringency, 2 pulldowns were performed per biological replicate. Streptavidin-coated magnetic beads (Pierce) were used to enrich for biotinylated proteins in a 96-well plate on an Agilent Bravo liquid-handler, using an in-house protocol adapted from Lam et. al.^53^. Briefly, 250 µg of cell lysate was incubated with 10 µL of equilibrated bead slurry for 90 minutes at 4°C. The beads were then washed twice with modified RIPA lysis buffer (without protease inhibitors), once with 1M KCL, twice more with lysis buffer and finally once with PBS. The beads were then resuspended in 50 mM ammonium bicarbonate (Sigma Aldrich) and the samples were reduced with 10 mM DTT (Thermo Fisher Scientific) for 30 minutes at room temperature. Following this, alkylation was performed in the dark with 55 mM 2-chloroacteamide (Sigma Aldrich), for 30-minutes at room temperature. The enriched proteins were then digested with trypsin (0.2 µg per sample, Promega) overnight at 37°C (with shaking at 1450 RPM) on a Thermomixer R (Eppendorf). The supernatants were collected with the help of a magnetic plate and topped up to 100 µL with 50 mM ammonium bicarbonate and 5 µL of 10% trifluoracetic acid (Thermo Fisher Scientific), and desalted using the StAGE-tipping protocol^54^ on the Bravo liquid handler with the AssayMap head and C18 peptide cleanup tips (Agilent). The peptides were dried using a SpeedVac (Eppendorf) concentrator and stored at -20°C.

#### Data-independent acquisition mass spectrometry and data analysis for turboID experiments

For the turboID experiments, the dried peptides were resuspended in reconstitution buffer (0.1% formic acid and 0.5% acetonitrile). 75 ng of peptides were injected and separated on-line using an Easy-nLC 1200 (Thermo Fisher Scientific) with Aurora Series analytical column (25cm x 75μm 1.6μm C18; Ion Opticks). The analytical column was heated to 40°C using an integrated column oven (PRSO-V2, Sonation). Buffer A consisted of 0.1% aqueous formic acid and 2% acetonitrile in water, and buffer B consisted of 0.1% aqueous formic acid and 80% acetonitrile in water. The analysis was performed at 0.25 μL/min flow rate with a standard 60min gradient: 1 min 2% B, 2% B to 20% B over 45 min, then to 32% B over 15 min, then to 50% B from 61 to 66 min, then to 95% B over 5 min, held at 95% B for 8 min, then dropped to 3% B over 2 min, held at 3% B for 6 min. The Easy-nLC thermostat temperature was set at 7°C. The peptides were analyzed with an Orbitrap Exploris 480 mass spectrometer (Orbitrap Exploris^TM^ 480, Thermo Fisher Scientific). The Nanospray Flex^TM^ ion source was operated at 1900 V spray voltage, and the ion transfer tube was heated to 290°C. During analysis, the Orbitrap Exploris 480 was operated in data-independent (DIA) mode in positive mode. Full scan MS resolution was set to 60,000 with 300% normalized automatic gain control (AGC) target, 50% RF lens, 25 ms maximum injection time and scan range from m/z 380 Th to m/z 985 Th. DIA fragment spectra were collected at a resolution of 15,000, normalized AGC target of 2000%, maximum injection time of 40 ms, scan range from m/z 145 Th – m/z 1450 Th. Isolation windows of m/z 10 Th (spanned the range from m/z 379.5 Th to m/z 980.5 Th) were used with an overlap of m/z 1 Th. Normalized collision energy was set to 28%. The mass-to-charge ratio was calibrated based on three selected ions from Pierce^TM^ FlexMix^TM^ Calibration Solution (m/z [Th]: 195, 322, 524, 622, 922, 1122, 1222, 1322, 1422, 1522, 1622, 1722 and 1822). The mass accuracy was typically within 2ppm and is not allowed to exceed 4ppm. For the first BioID experiment, each sample was loaded twice onto the instrument for additional technical replicates, while for the second one, samples were loaded once.

The RAW files were analyzed with DIA-NN 1.9.2^55^, and a human FASTA database from Uniprot was used to generate an *in-silico* spectral library. The FASTA database was supplemented with sequences for streptavidin (UniprotID – P22629), trypsin (UniprotID – P00761), *Renilla* luciferase (UniprotID – P27652) and the turboID sequence. The raw fragment quantities were also exported by including the “--export-quant” command in the additional options section. The DIA-NN output parquet file was filtered for Q.Value < 0.01, PG.Q.Value < 0.05, Lib.Q.Value < 0.01, Lib.PG.Q.Value < 0.01. This was done to exclude low confidence proteins and precursors. To score true protein interactors, the fragment-level analysis offered by SAINTq 0.0.4^56^ was used. The RLuc samples were set as the negative control, and fragment intensities were median normalized prior to SAINTq analysis. For the first BioID experiment, the peptide selection rules were set to min_n_pep = 3, best_prop_pep = 0.35 and fragment selection rules were set to min_n_frag = 3, best_prop_frag = 0.35. Test and control baits were set at compress_n_rep=4. For the second BioID experiment, the peptide selection rules were set to min_n_pep = 3,best_prop_pep = 0.45 and fragment selection rules were set to min_n_frag = 3, best_prop_frag = 0.45. Test and control baits were set at compress_n_rep=2. Following the identification of high confidence, wildtype and mutant interactors, a 2-sample, unpaired Welch’s t-test was used to identify proteins that were enriched in mutant samples (log2 FC >= 1.0 and Benjamini-Hochberg corrected P-values < 0.05). For this, the PG.MaxLFQ values from the filtered DIA-NN report were used as protein intensities. This intensity matrix was median normalized and missing values were imputed using the imputeLCMD package in R (q=0.01 and sigma = 0.005).

#### siRNA transfections to establish functional relationships

For the siRNA experiments performed in Figure 2 and 3C, the silencer select siRNA from Thermo Fisher was used (all RNAi oligos are listed in Table S4). 15 nM or 30 nM of each siRNA was reverse transfected into the indicated cell lines using Lipofectamine™ RNAiMAX (Invitrogen) using the manufacturer’s protocol. These were performed in 96-well plates, and the knockdown was carried out over a period of 72-hours. For the BAG6 and RNF126 knockdown experiments shown in Figure 3B, the 12.5 nM of ON-TARGETplus siRNA – SMARTpool from Dharmacon were reverse transfected into cells grown in 6-well plates (35 mm) using Lipofectamine™ RNAiMAX. Knockdowns were carried out over a period of 72-hours. For experiments shown in Figure 5F cells were reverse transfected with ON-TARGETplus siRNA – SMARTpool using Dharmafect (Horizon Discovery) according to the recommended protocol. All siRNA sequences, and catalog numbers have been provided in Table 4. For experiments performed in Figures 2 and 3, cells grown in 96- or 6-well plates were trypsinized 72-hours post-siRNA transfection and transferred to 96-well v-bottom plates (Sarstedt). For cells grown in 96-well plates, all cells were processed for flow cytometry, while for cells grown in 6-well plates, half the cells were processed for flow cytometry, and the other half were lysed for a Western blot analysis to assess protein knockdown. The cells were pelleted by centrifugation at 400*g* (5 minutes at 4°C), washed once with cold PBS and then fixed with 2% formaldehyde (15 minutes at room temperature). The samples were then washed twice with cool PBS and then analyzed on the Cytoflex (Beckman) and at least 20,000 events were collected. For flow cytometry experiments shown in Figures 4 and 5, live cells (suspended in 2% FBS in PBS) were analyzed and 30,000 events per sample were collected. Cells were gated on the FSC-A v/s SSC-A, and FSC-A v/s FSC-W parameters. Cells expressing the reporter were isolating by gating cells that showed > 10^4^ arbitrary fluorescence units in the ECD-A channel.

#### Flow cytometry experiments to assess degron stability

For exogenous expression of degrons, the constructs were transfected into HEK293T or HepG2 cells using FuGENE (Promega). 48 hours (HEK293T) or 72 hours (HepG2) after transfection, live cells were harvested and analyzed on Cytoflex (Beckman) as described above. For HEK293T cells, 50,000 events were collected, while for HepG2 cells, 100,000 events were collected. Analysis and gating strategy were identical to the one described above. For stable cell lines, data were collected using a similar approach.

#### Western blot and dot blot analysis

Cells were lysed in modified RIPA buffer as described above and lysates were diluted in Laemmli sample buffer and boiled for 5 minutes. The samples were resolved by SDS-PAGE on 4-20% TGX gradient gels (BioRad) and transferred to 0.45 mm nitrocellulose membranes using a Trans-Blot Turbo Transfer System at 25 V for 30 min (Bio-Rad). The Revert™ 700 Total Protein Stain (Licor) was used following the manufacturer’s protocol and visualized on the Odyssey Imager (Licor). Membranes were blocked with 5% non-fat milk in TBS-T and probed with primary antibodies overnight at 4°C (all antibodies used in this study are listed in Table S5). After washing with 1X TBS-T (3x 5 min), membranes were incubated secondary antibodies (conjugated with either IR Dye 680RD or 800CW, Licor) and washed again. Bands were visualized using the Licor Odyssey imager. For dot blotting, 3 µL of denatured samples in Laemmli sample buffer were directly spotted onto 0.45 mm nitrocellulose membranes, following which the aforementioned Western blotting protocol was followed. For capillary analysis in Figure S1, the Jess system (ProteinSimple) was utilized, following the manufacturer’s protocol.

#### Data-independent mass spectrometry and data analysis for siRNA validation

To validate siRNA knockdowns shown in Figure 3C, siRNA transfected cells were lysed in modified RIPA buffer as described above. 10 µg of protein were processed with Sera-Mag™ carboxylate-modified SpeedBeads (Cytiva), following the previously published SP3 protocol^57^. The proteins were digested using sequencing grade trypsin (Promega) and then desalted using the aforementioned StAGE-tipping protocol. 75 ng of peptides were then loaded onto the Exploris 480 for DIA-mass spectrometry as described above. Data was analyzed using DIA-NN 2.0.2.

#### Degron predictor

The support vector regression models were trained on GPS-peptidome data in generated by Zhang et al^25^ containing ∼260K unique peptide sequences. C-terminal degron positive controls were excluded from the training dataset to avoid detection of C-terminal degrons for protein-internal sequences, leaving behind a 252K internal degron dataset. Features for each peptide sequence were first extracted in Python from the peptide sequence and the feature output .csv used as input for the support vector regression model in R. Both programs and supporting data are available at (https://github.com/Alpine2048/Degron-prediction-by-composition-seq-motifs). Models were trained on 1% of the filtered dataset (2525 peptides) using the svmRadial method provided by the Caret package for R and ranked by testing on the remaining 99% of peptide sequence. The CSM* model that includes the experiment specific flanking residues was used for the GNMT degron prediction calculations.

#### AlphaFold3 predictions

AlphaFold version 3.0.1^58^ was used to predict binary protein-protein interactions between GNMT degron (residues 281-295) and the defined domains of Bag6, using default parameters. For each prediction, 5 models were generated, and the highest-scoring model was retained for further analysis. High-scoring models were further assessed using predicted template modelling (pTM), interface-pTM (ipTM), and combined score (calculated as 0.8*ipTM + 0.2*pTM)^59^. AlphaFold3 predictions were carried out on an Nvidia H100 GPU within the Digital Research Alliance of Canada Fir cluster and reported values are listed in Table S7.

#### Data analysis and figure generation

Data analysis and statistics were performed in Microsoft Excel, R (RStudio version 2023.06.0; R version 4.4.2) and GraphPad Prism. Heat maps shown in Figures 1,2 and S3 were generated with the pheatmap package. The upset plot shown in Figure S3 generated using the upsetR package. Gene ontology analyses were performed with DAVID^60,61^. All figures were generated in Illustrator.

## Supplemental information

Document S1: Figures S1–S6, Tables S3, S4, S5, S6 and S7

Supplemental Table 1 & 2 (MS data for first and second turboID experiments)

---

**Figure S1.**
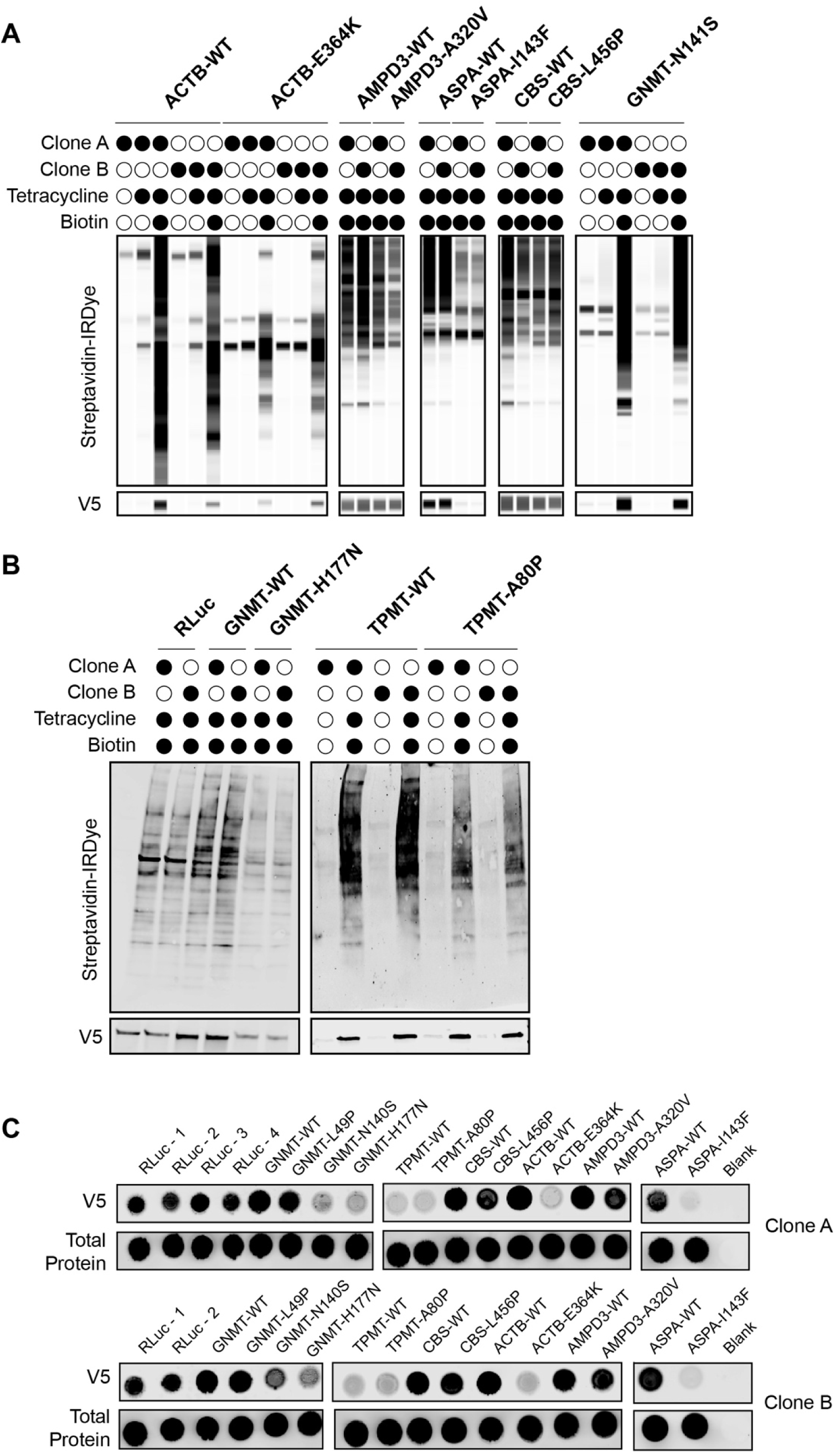
BioID baits. **A.** Jess analysis of the indicated BioID baits. **B.** Western blot analysis of the indicated remaining baits. **C.** Dot blot analysis of the indicated samples that were processed for the BioID pull downs (wildtype and mutant samples were processed on the same blot).

**Figure S2.**
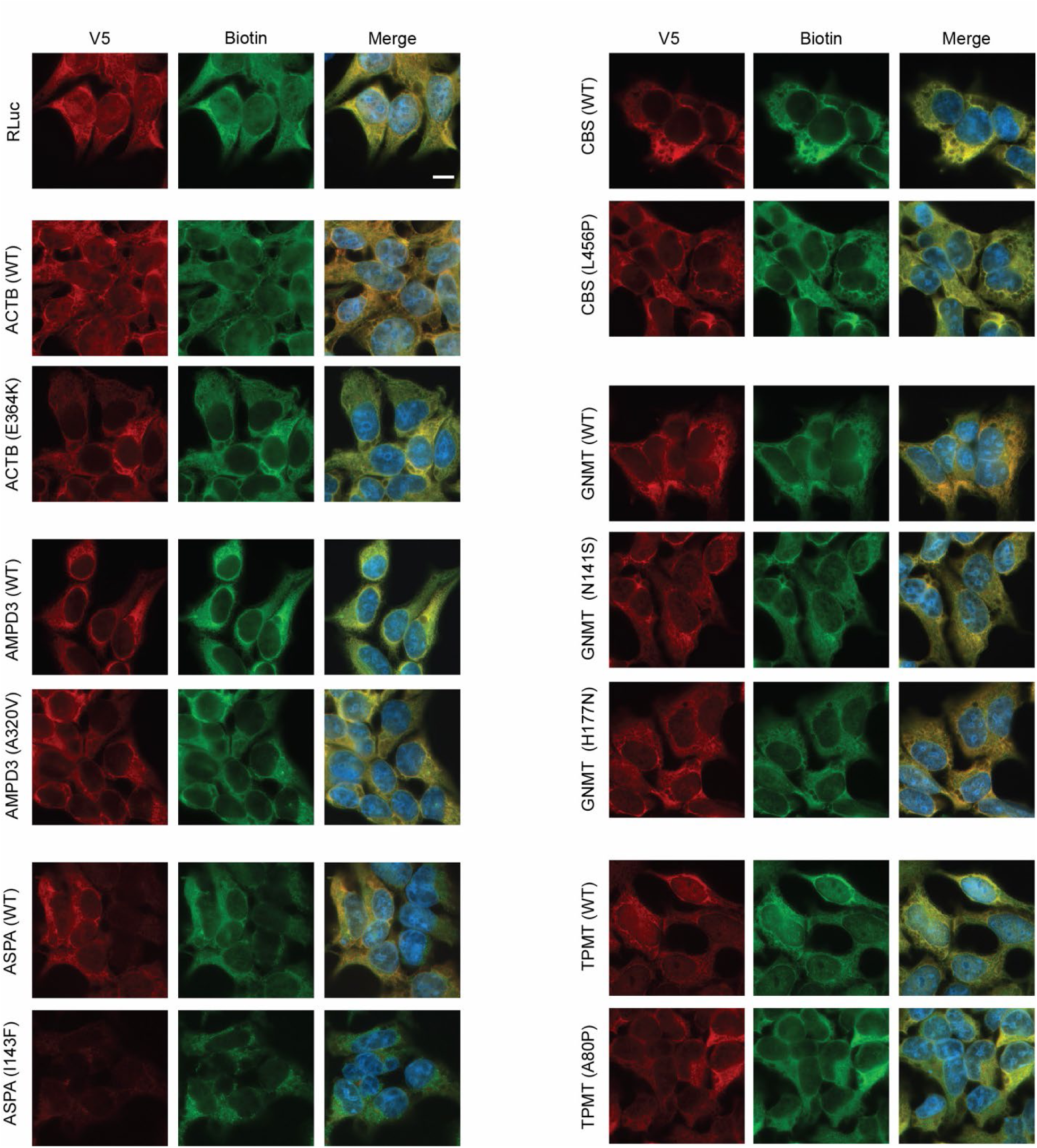
Imaging of BioID baits. Immunofluorescence microscopy of HEK293 cells expressing the indicated baits for 24 hours and treated with biotin for 30 minutes. Exposure settings were adjusted to facilitate visualization of mutant variants expressed at lower levels. Scale bar, 10 µm.

**Figure S3.**
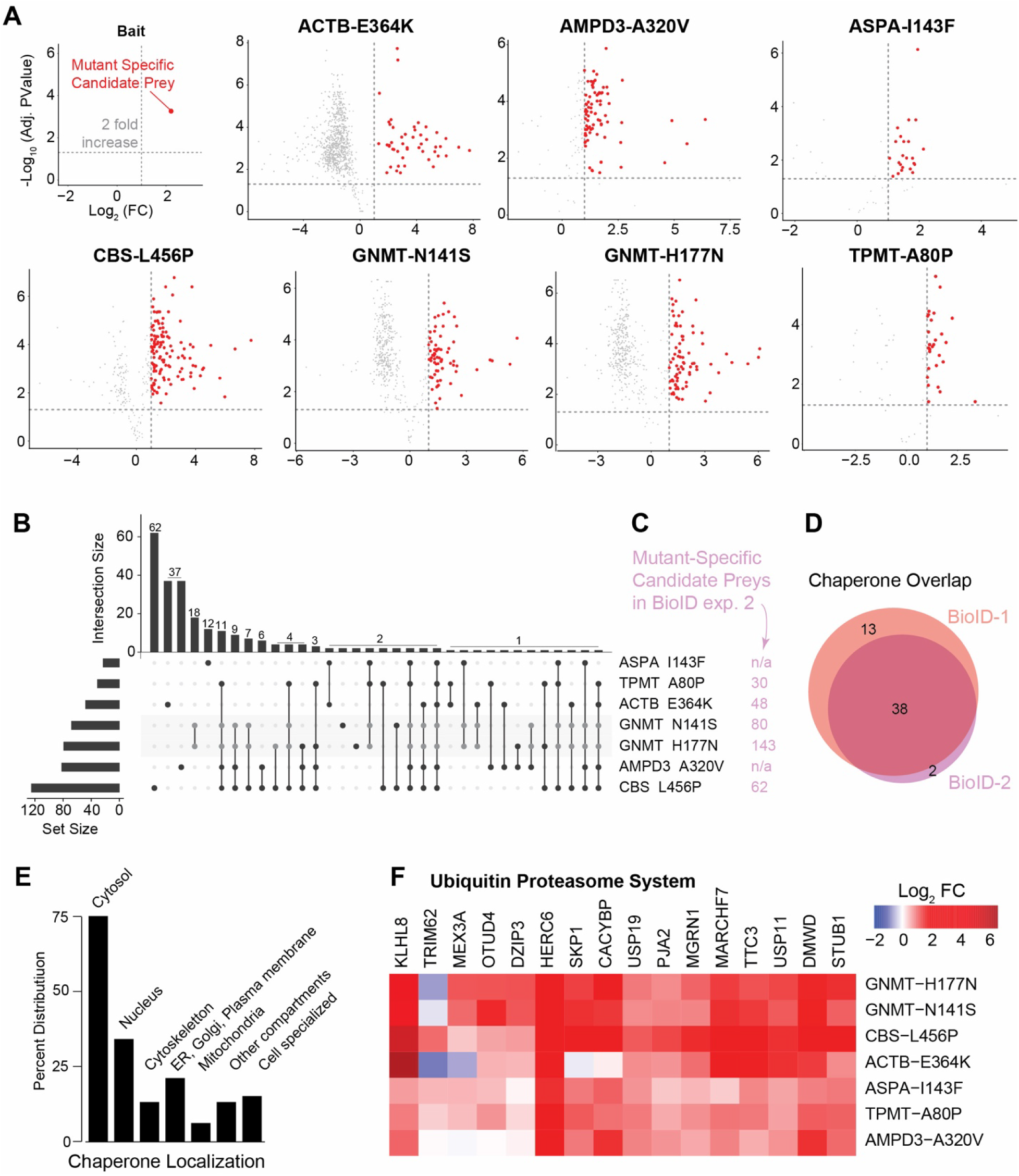
BioID experiment. **A.** Volcano plots showing the log₂ fold change in signal intensity of high-confidence candidate interactors in mutant pulldowns compared to their wild-type counterparts. Proteins with a ≥2-fold change and adjusted p-value < 0.05 were considered mutant-specific preys (depicted in red; n = 2 technical replicates for 2 biological replicates). **B.** UpSet plot of mutant-specific preys for the indicated variants. **C.** Number of preys identified in an independent BioID experiment in which ASPA and AMPD3 were not assessed (n = 2 technical replicates for 2 biological replicates). **D.** Overlap of chaperone proteins identified as mutant-specific preys across both BioID experiments. **E.** Subcellular localization of 46 chaperones with available annotations in Protein Atlas. Some proteins have multiple reported localizations. **F.** Heat map of mutant-specific preys involved in ubiquitination, as shown in Figure 1D.

**Figure S4.**
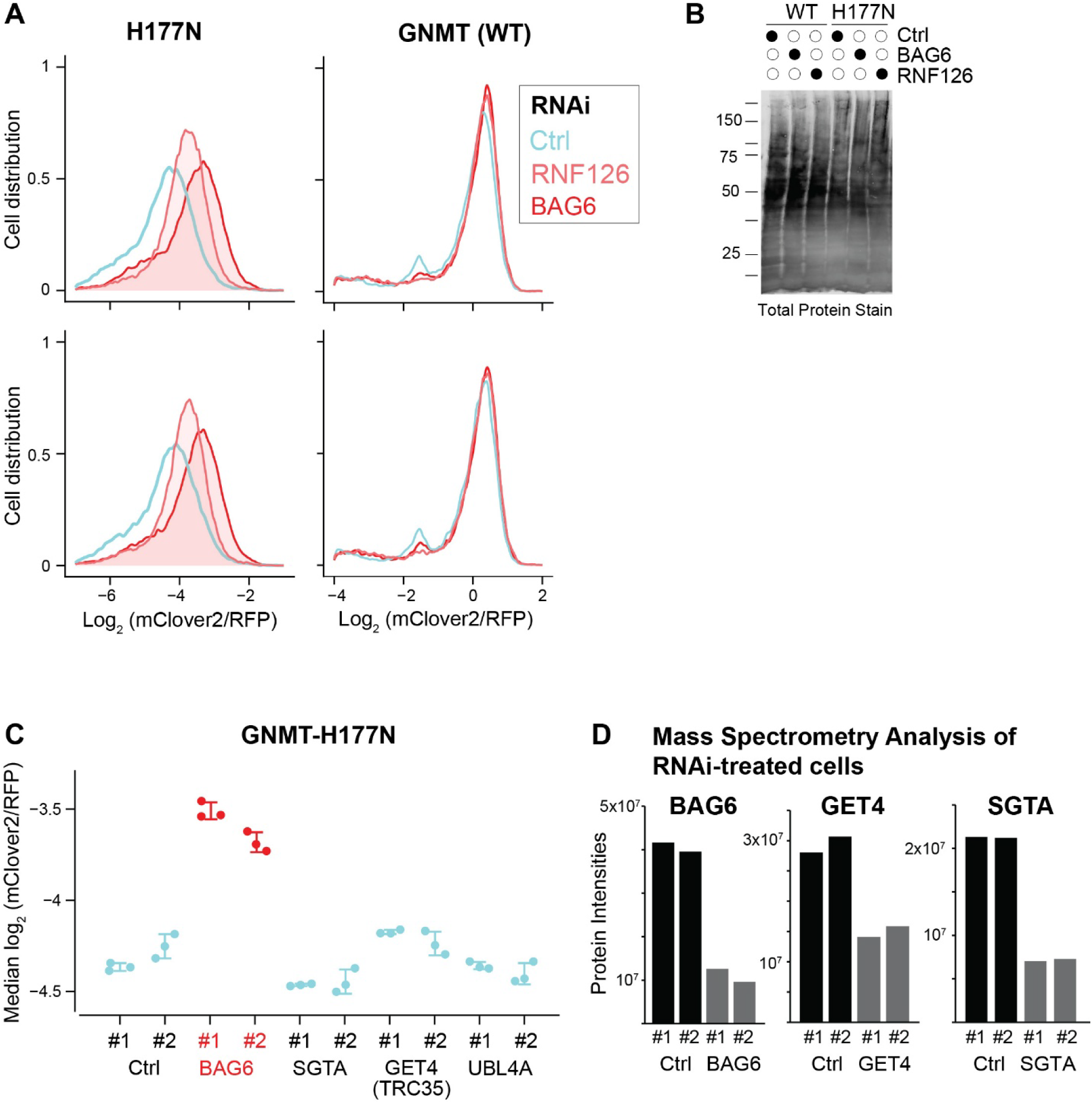
Involvement of BAG6 and RNF126 in GNMT mutant degradation. **A.** Flow cytometry analysis of cells transduced with WT or mutant GNMT reporters and transfected with the indicated RNAi. These data represent two additional experiments complementing the analysis shown in Figure 3B. **B.** Total protein staining of the blot presented in Figure 3B. **C.** Median log₂ ratio of mClover to RFP signal for GNMT H177N expressed in HEK293 cells (n = 3 for each indicated RNAi). The average of these median values is shown in Figure 3C. **D.** Intensities of the indicated proteins quantified by mass spectrometry of total cell extracts of the indicated samples.

**Figure S5.**
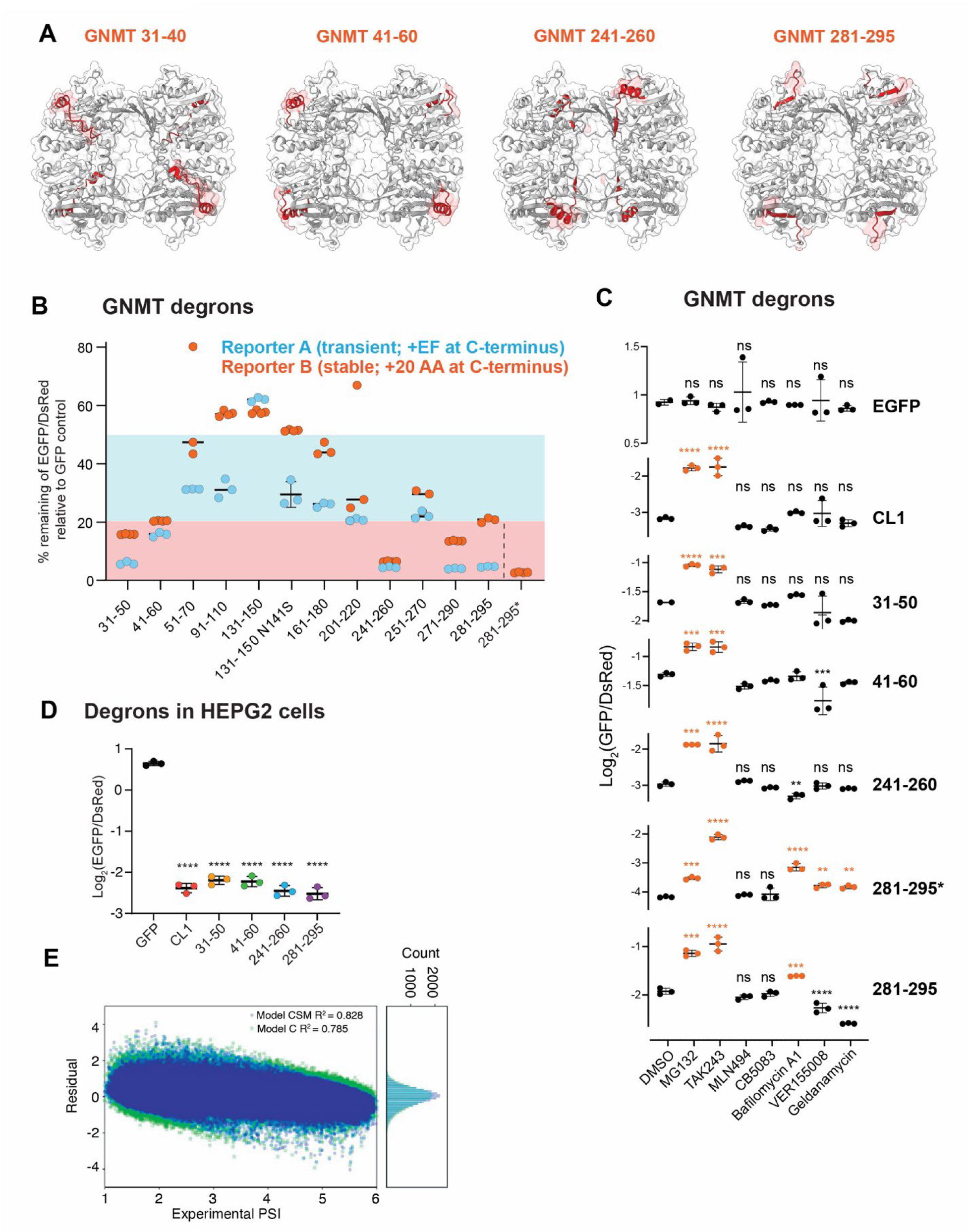
Identification and characterization of GNMT degrons. **A.** Mapping of the indicated degrons onto the GNMT tetramer structure using ChimeraX^64–66^ (1R74). **B.** Normalized levels of the indicated GNMT tiles, either transfected with the reporter containing the EF C-terminal region (same data as in Figure 4B) or transduced with the reporter containing the 20–amino acid C-terminal region (n = 3; * indicates a tile containing a stop codon following the last GNMT residue). **C-D.** Detailed data corresponding to Figure 4C (n = 3) and Figure 4D (n = 3; unpaired Student’s t-test, ns, *, **, ***, and **** correspond to p > 0.05, < 0.05, < 0.01, < 0.001, and < 0.0001, respectively). **E.** Plot of residuals (PSI_experimental_ – PSI_predicted_) versus experimental PSI (Protein Stability Index) for two models: Model C (amino acid fractional composition only) and CSM (Model C with sequence motifs). The R² values for predicted versus measured PSI and adjacent histograms of residuals are shown.

**Figure S6.**
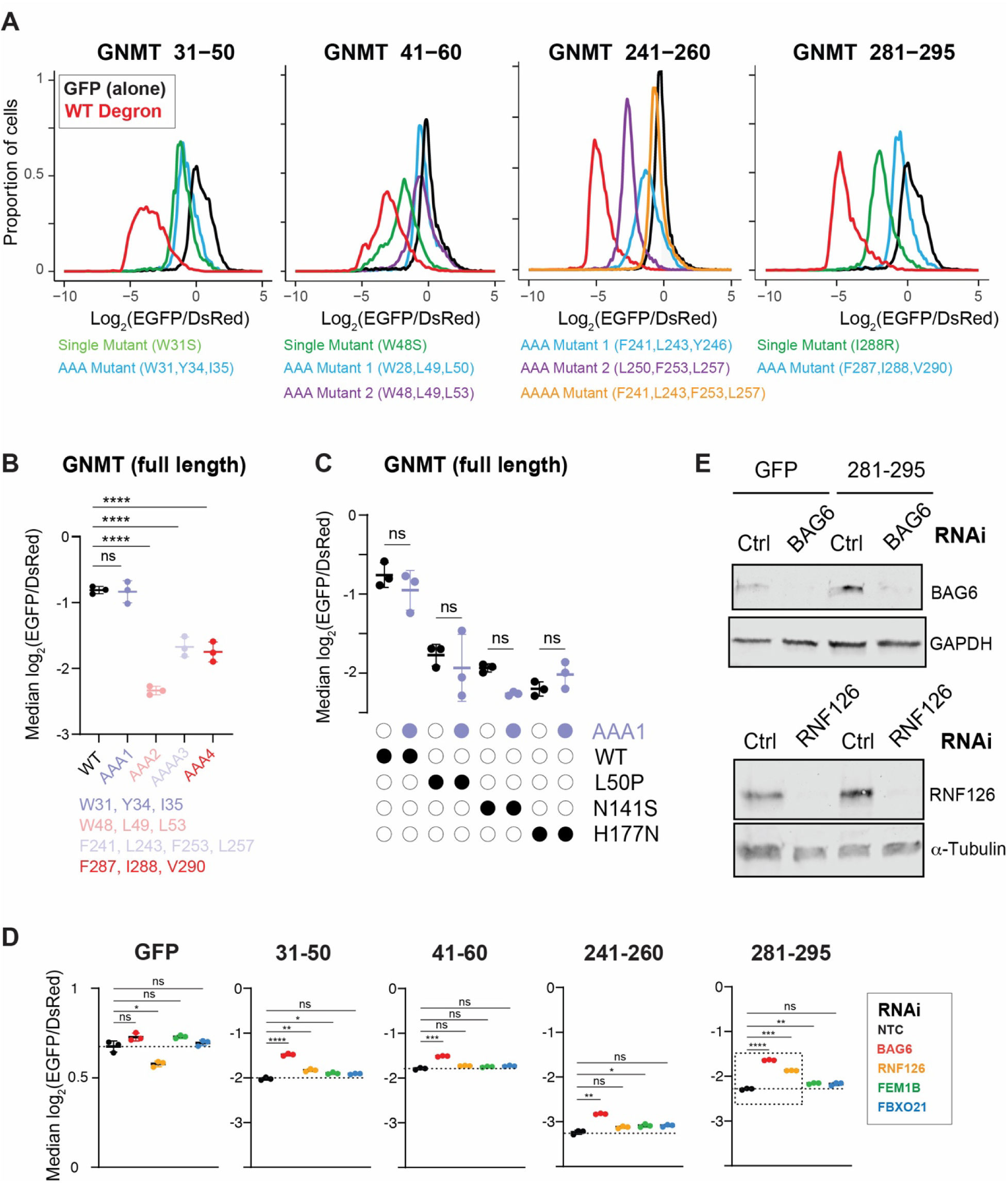
Identification of GNMT degron sequences and impact of RNAi on degradation. **A.** Flow cytometry analysis of the indicated mutations in the assessed tiles (n = 1). **B.** Flow cytometry analysis of the indicated mutations in full-length GNMT (n = 3; unpaired t-test; ns, p > 0.05; ****, p < 0.0001). **C.** Flow cytometry analysis of W31A, Y34A, and I35A mutations in the indicated full-length GNMT variants (n = 3; unpaired t-test; ns, p > 0.05). **D.** Flow cytometry analysis of the indicated tiles expressed in HEK293 cells following the specified RNAi treatments (n = 3; unpaired t-test; ns, *, **, ***, and **** correspond to p > 0.05, < 0.05, < 0.01, < 0.001, and < 0.0001, respectively).

