## Supplemental Figures and Tables for "BAG6 and RNF126 promote the degradation of cytosolic misfolded proteins that contain buried degron motifs"

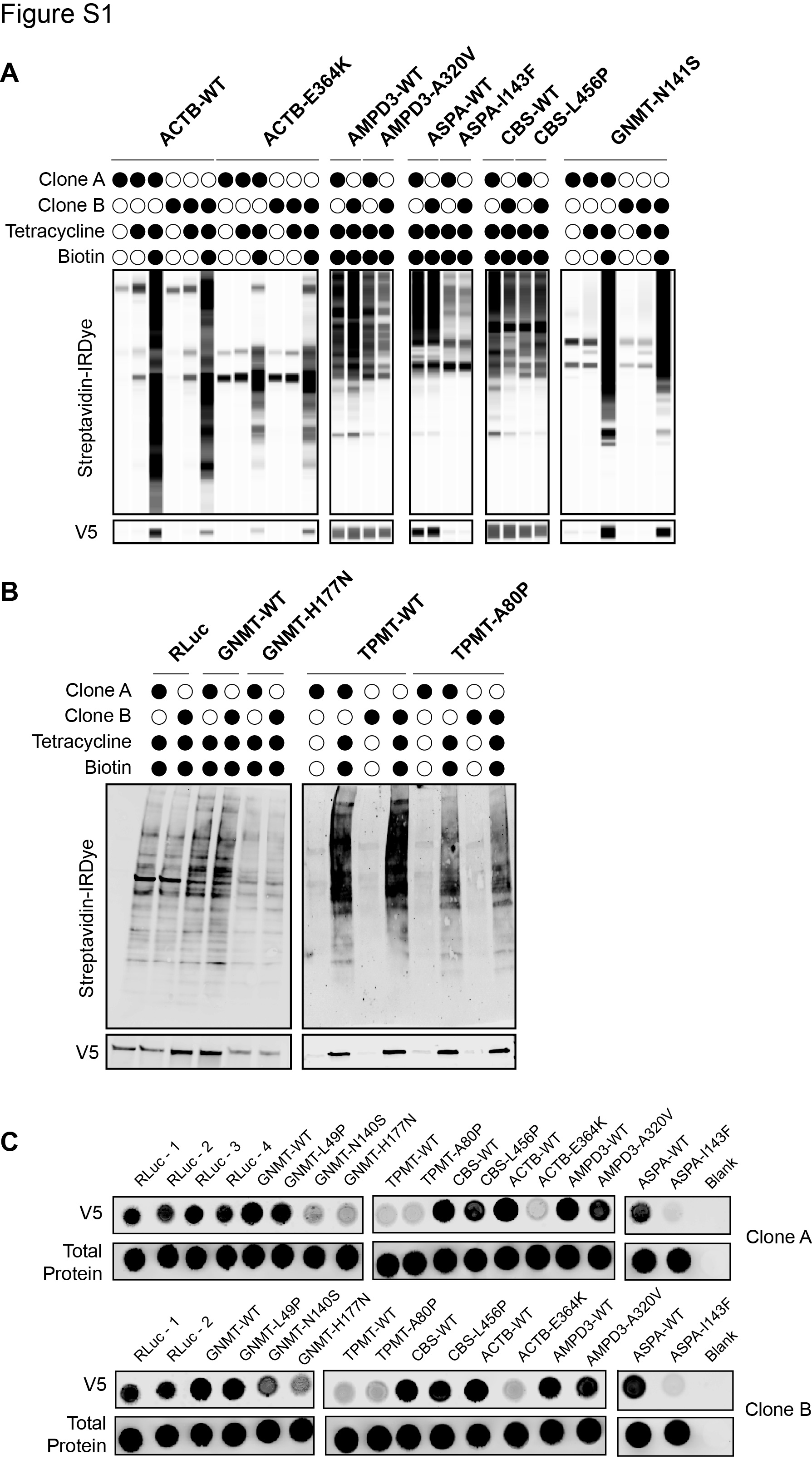

**Figure S1. BioID baits. A.** Jess analysis of the indicated BioID baits. **B.** Western blot analysis of the indicated remaining baits. **C.** Dot blot analysis of the indicated samples that were processed for the BioID pull downs (wildtype and mutant were processed on the same blot).

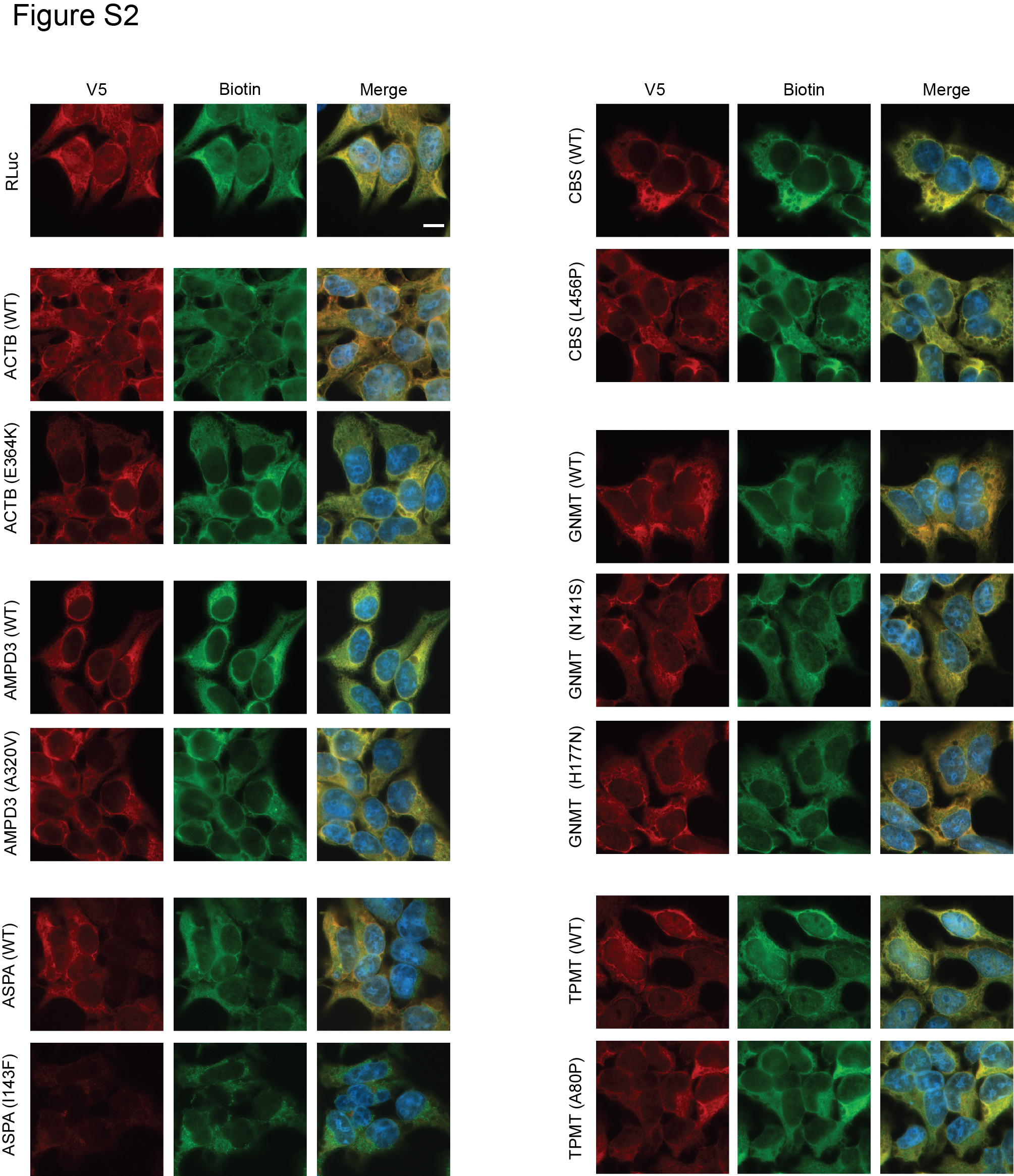

**Figure S2. Imaging of BioID baits.** Immunofluorescence microscopy of HEK293 cells expressing the indicated baits for 24 hours and treated with biotin for 30 minutes. Exposure settings were adjusted to facilitate visualization of mutant variants expressed at lower levels. Scale bar, 10 µm.

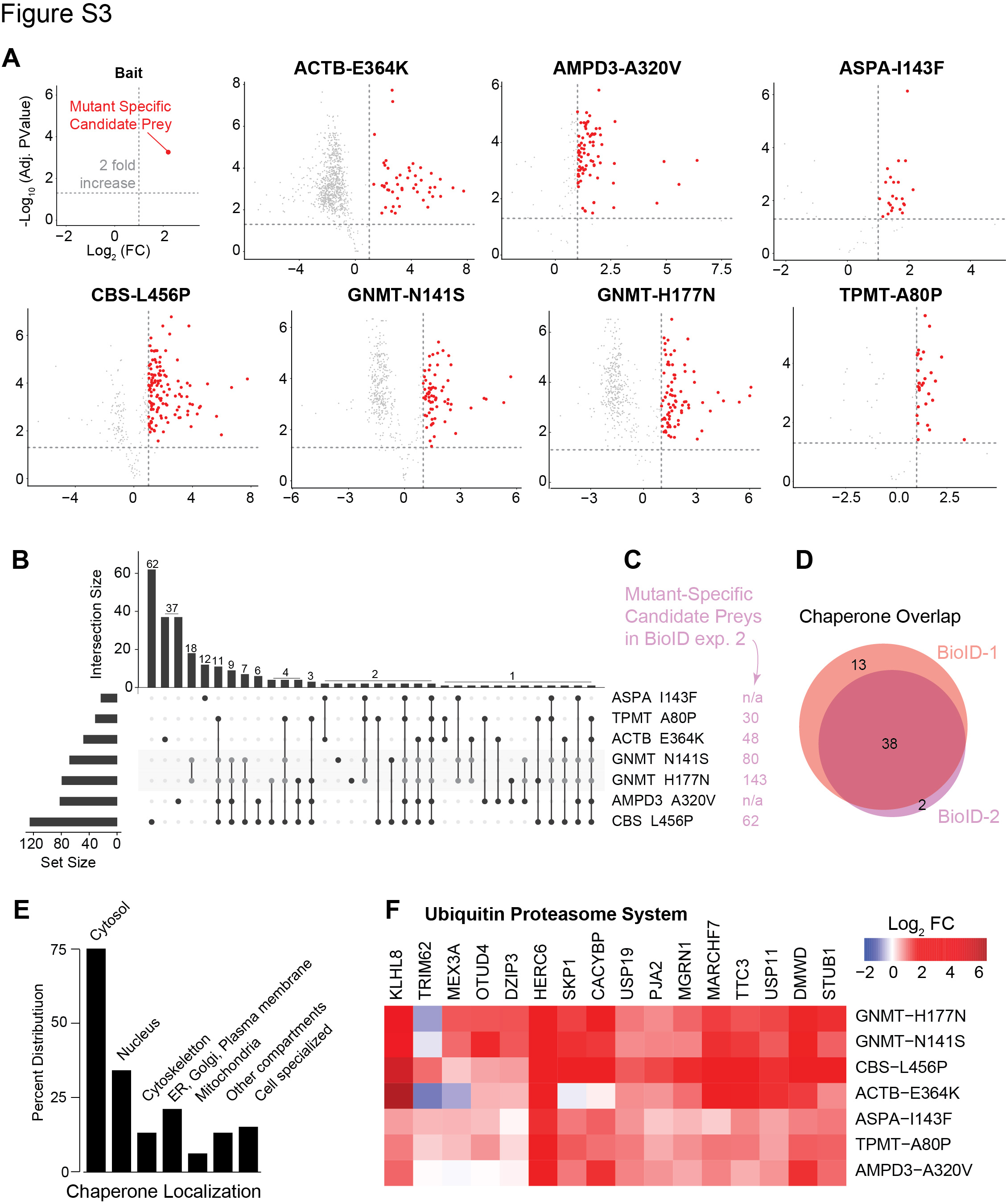

**Figure S3. BioID experiment.** **A.** Volcano plots showing the log₂ fold change in signal intensity of high-confidence candidate interactors in mutant pulldowns compared to their wild-type counterparts. Proteins with a ≥2-fold change and adjusted p-value < 0.05 were considered mutant-specific preys (depicted in red; n = 2 technical replicates for 2 biological replicates). **B.** UpSet plot of mutant-specific preys for the indicated variants. **C.** Number of preys identified in an independent BioID experiment in which ASPA and AMPD3 were not assessed (n = 2 technical replicates for 2 biological replicates). **D.** Overlap of chaperone proteins identified as mutant-specific preys across both BioID experiments. **E.** Subcellular localization of 46 chaperones with available annotations in Protein Atlas. Some proteins have multiple reported localizations. **F.** Heat map of mutant-specific preys involved in ubiquitination, as shown in Figure 1D.

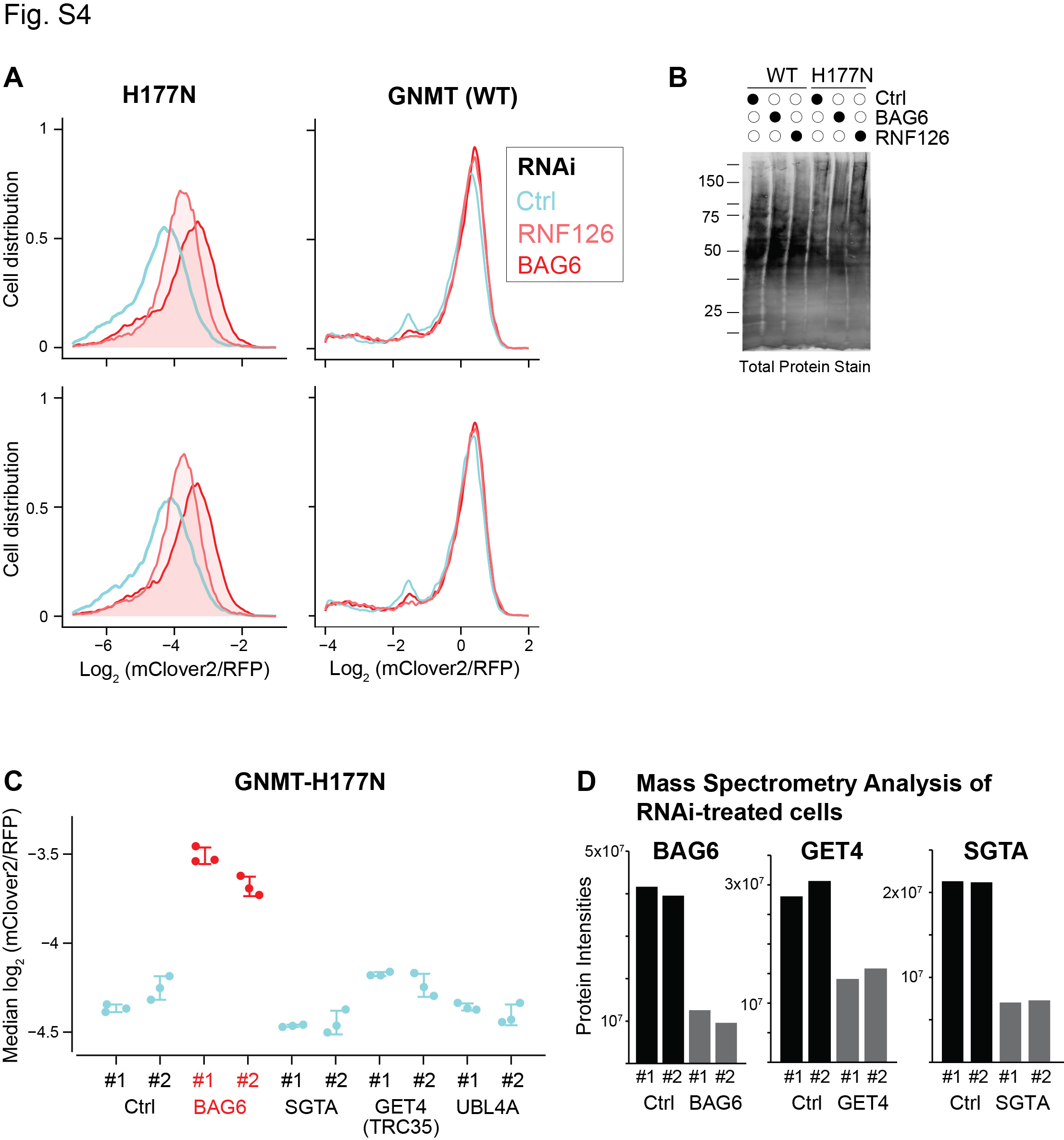

**Figure S4. Involvement of BAG6 and RNF126 in GNMT mutant degradation.** **A.** Flow cytometry analysis of cells transduced with WT or mutant GNMT reporters and transfected with the indicated RNAi. These data represent two additional experiments complementing the analysis shown in Figure 3B. **B.** Total protein staining of the blot presented in Figure 3B. **C.** Median log₂ ratio of mClover to RFP signal for GNMT H177N expressed in HEK293 cells (n = 3 for each indicated RNAi). The average of these median values is shown in Figure 3C. **D.** Intensities of the indicated proteins quantified by mass spectrometry of total cell extracts of the indicated samples.

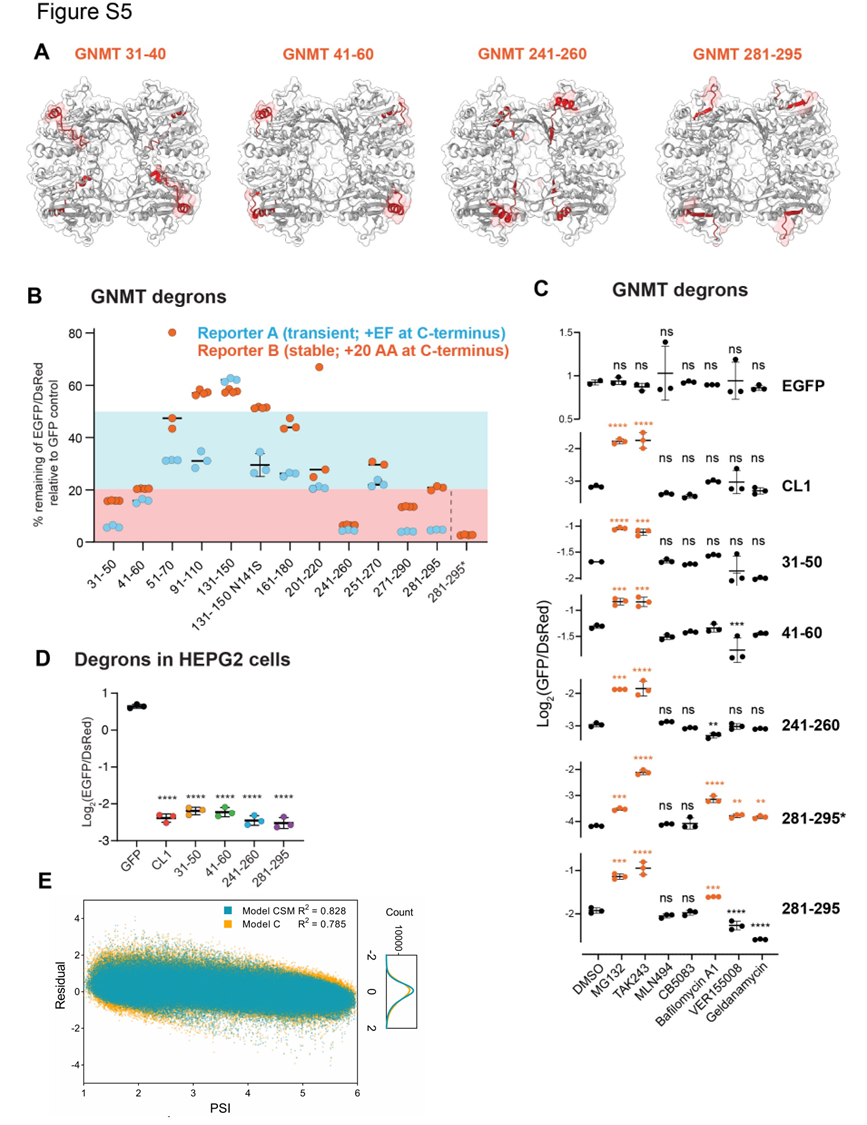

**Figure S5. Identification and characterization of GNMT degrons. A.** Mapping of the indicated degrons onto the GNMT tetramer structure using ChimeraX (1R74). **B.** Normalized levels of the indicated GNMT tiles, either transfected with the reporter containing the EF C-terminal region (same data as in Figure 4B) or transduced with the reporter containing the 20–amino acid C-terminal region (n = 3; * indicates a tile containing a stop codon following the last GNMT residue). **C-D.** Detailed data corresponding to Figure 4C (n = 3) and Figure 4D (n = 3; unpaired Student’s t-test, ns, *, **, ***, and **** correspond to p > 0.05, < 0.05, < 0.01, < 0.001, and < 0.0001, respectively). **E.** Plot of residuals (PSI_experimental_ – PSI_predicted_) versus experimental PSI (Protein Stability Index) for two models: Model C (amino acid fractional composition only) and CSM (Model C with sequence motifs). The R² values for predicted versus measured PSI and adjacent histograms of residuals are shown.

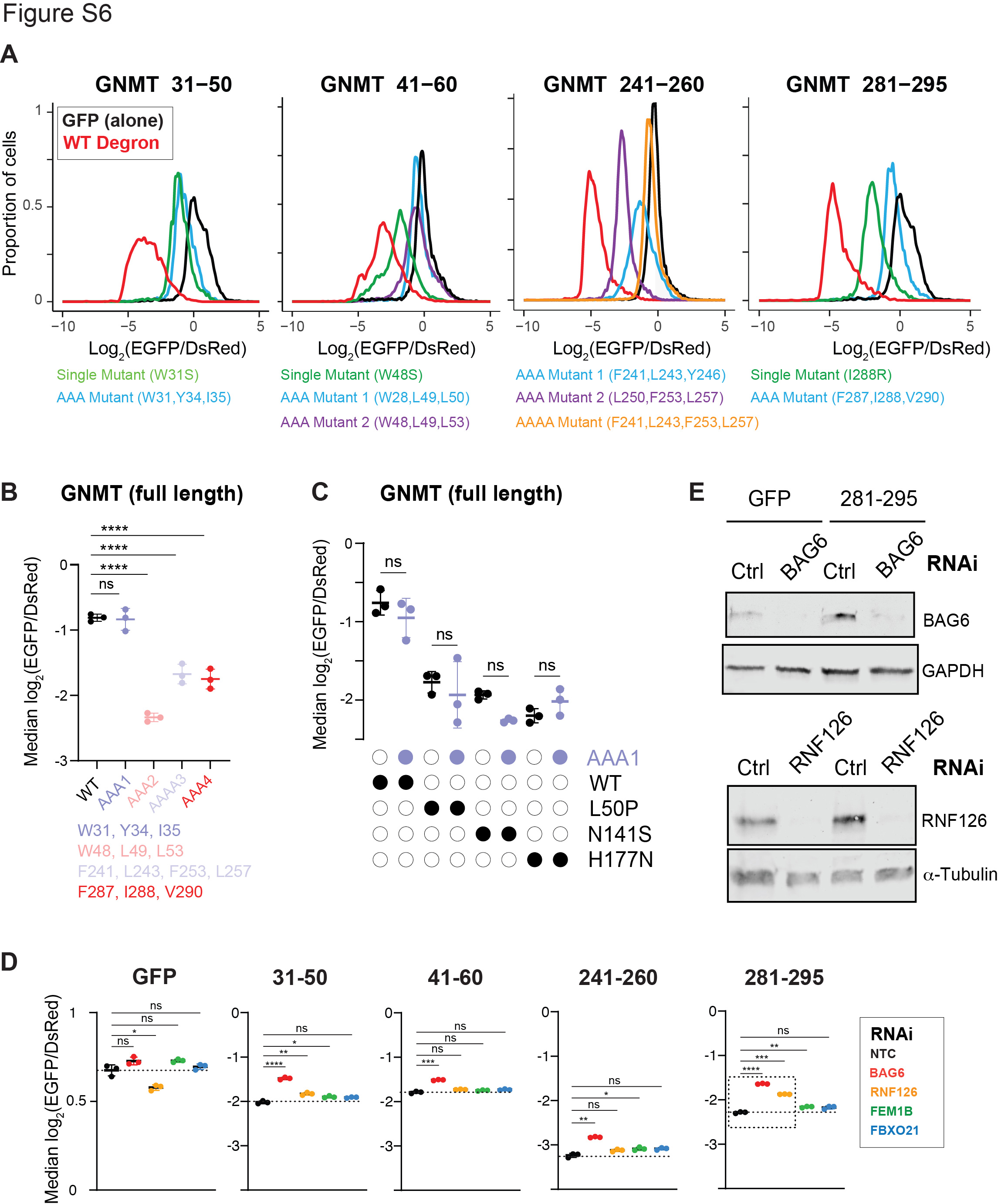

**Figure S6. Identification of GNMT degron sequences and impact of RNAi on degradation.** **A.** Flow cytometry analysis of the indicated mutations in the assessed tiles (n = 1). **B.** Flow cytometry analysis of the indicated mutations in full-length GNMT (n = 3; unpaired t-test; ns, p > 0.05; ****, p < 0.0001). **C.** Flow cytometry analysis of W31A, Y34A, and I35A mutations in the indicated full-length GNMT variants (n = 3; unpaired t-test; ns, p > 0.05). **D.** Flow cytometry analysis of the indicated tiles expressed in HEK293 cells following the specified RNAi treatments (n = 3; unpaired t-test; ns, *, **, ***, and **** correspond to p > 0.05, < 0.05, < 0.01, < 0.001, and < 0.0001, respectively).

**Supplementary Table S1:**

Excel

**Supplementary Table S2:**

Excel

**Supplementary Table 3: Plasmids used in this study**

| **Plasmid** | **Backbone** | **Gene** | **Mayor Lab BPM** |
| --- | --- | --- | --- |
| TPMT-A80P-V5-TurboID | pcDNA5 FRT/TO | TPMT-A80P | 1944 |
| TPMT-V5-TurboID | pcDNA5 FRT/TO | TPMT | 1943 |
| GNMT H177N-V5-TurboID | pcDNA5 FRT/TO | GNMT-H177N | 1920 |
| GNMT-V5-TurboID | pcDNA5 FRT/TO | GNMT | 1919 |
| GNMT-N141S-V5-TurboID | pcDNA5 FRT/TO | GNMT-N141S | 1947 |
| AMPD3-V5-TurboID | pcDNA5 FRT/TO | AMPD3 | 1949 |
| AMPD3-A320V-V5-TurboID | pcDNA5 FRT/TO | AMPD3-A320V | 1950 |
| ACTB-V5-TurboID | pcDNA5 FRT/TO | ACTB | 1955 |
| ACTB-E364K-V5-TurboID | pcDNA5 FRT/TO | ACTB-E364K | 1956 |
| ASPA-I143F-V5-TurboID | pcDNA5 FRT/TO | ASPA-I143F | 1957 |
| ASPA-V5-TurboID | pcDNA5 FRT/TO | ASPA | 1960 |
| CBS-V5-TurboID | pcDNA5 FRT/TO | CBS | 1958 |
| CBS-L456P-V5-TurboID | pcDNA5 FRT/TO | CBS-L456P | 1961 |
| RLuc-V5-TurboID | pcDNA5 FRT/TO | RLuc | 1876 |
| ACTB-E364K-mClover2-P2A-mRFP1 | pLenti6.2-ORF-mClover2-P2A-mRFP1 | ACTB-E364K | 1551 |
| CBS-L456P-mClover2-P2A-mRFP1 | pLenti6.2-ORF-mClover2-P2A-mRFP1 | CBS-L456P | 1553 |
| GNMT-mClover2-P2A-mRFP1 | pLenti6.2-ORF-mClover2-P2A-mRFP1 | GNMT | 1557 |
| GNMT H177N-mClover2-P2A-mRFP1 | pLenti6.2-ORF-mClover2-P2A-mRFP1 | GNMT-H177N | 1558 |
| TPMT A80P-mClover2-P2A-mRFP1 | pLenti6.2-ORF-mClover2-P2A-mRFP1 | TPMT-A80P | 1626 |
| CMV-DsRed-IRES-EGFP-GNMT 281-295 AAA | MSCV | GNMT 281-295 AAA | 2141 |
| CMV-DsRed-IRES-EGFP-GNMT 271-290 AAA | MSCV | GNMT 271-290 AAA | 2140 |
| CMV-DsRed-IRES-EGFP-GNMT 241-260 AAA | MSCV | GNMT 241-260 AAA | 2138 |
| CMV-DsRed-IRES-EGFP-GNMT 41-60 AAA | MSCV | GNMT 41-60 AAA | 2136 |
| CMV-DsRed-IRES-EGFP-GNMT 31-50 AAA | MSCV | GNMT 31-50 AAA | 2135 |
| CMV-DsRed-IRES-EGFP-GNMT 151-170 | MSCV | GNMT 151-170 | 2000 |
| CMV-DsRed-IRES-EGFP-GNMT 261-280 | MSCV | GNMT 261-280 | 1999 |
| CMV-DsRed-IRES-EGFP-GNMT 251-270 | MSCV | GNMT 251-270 | 1998 |
| CMV-DsRed-IRES-EGFP-GNMT 241-260 | MSCV | GNMT 241-260 | 1997 |
| CMV-DsRed-IRES-EGFP-GNMT 231-250 | MSCV | GNMT 231-250 | 1996 |
| CMV-DsRed-IRES-EGFP-GNMT 211-230 | MSCV | GNMT 211-230 | 1995 |
| CMV-DsRed-IRES-EGFP-GNMT 171-190 | MSCV | GNMT 171-190 | 1994 |
| CMV-DsRed-IRES-EGFP-GNMT 161-180 | MSCV | GNMT 161-180 | 1993 |
| CMV-DsRed-IRES-EGFP-GNMT 281-296 | MSCV | GNMT 281-296 | 1992 |
| CMV-DsRed-IRES-EGFP-GNMT 271-290 | MSCV | GNMT 271-290 | 1991 |
| CMV-DsRed-IRES-EGFP-GNMT 221-240 | MSCV | GNMT 221-240 | 1990 |
| CMV-DsRed-IRES-EGFP-GNMT 201-220 | MSCV | GNMT 201-220 | 1989 |
| CMV-DsRed-IRES-EGFP-GNMT 191-210 | MSCV | GNMT 191-210 | 1988 |
| CMV-DsRed-IRES-EGFP-GNMT 181-200 | MSCV | GNMT 181-200 | 1987 |
| CMV-DsRed-IRES-EGFP-GNMT 171-190 H177N | MSCV | GNMT 171-190 H177N | 1986 |
| CMV-DsRed-IRES-EGFP-GNMT 141-160 | MSCV | GNMT 141-160 | 1985 |
| CMV-DsRed-IRES-EGFP-GNMT 131-150 N141S | MSCV | GNMT 131-150 N141S | 1984 |
| CMV-DsRed-IRES-EGFP-GNMT 131-150 | MSCV | GNMT 131-150 | 1983 |
| CMV-DsRed-IRES-EGFP-GNMT 121-140 | MSCV | GNMT 121-140 | 1982 |
| CMV-DsRed-IRES-EGFP-GNMT 91-110 | MSCV | GNMT 91-110 | 1981 |
| CMV-DsRed-IRES-EGFP-GNMT 111-130 | MSCV | GNMT 111-130 | 1980 |
| CMV-DsRed-IRES-EGFP-GNMT 101-120 | MSCV | GNMT 101-120 | 1979 |
| CMV-DsRed-IRES-EGFP-GNMT 81-100 | MSCV | GNMT 81-100 | 1978 |
| CMV-DsRed-IRES-EGFP-GNMT 71-90 | MSCV | GNMT 71-90 | 1977 |
| CMV-DsRed-IRES-EGFP-GNMT 61-80 | MSCV | GNMT 61-80 | 1976 |
| CMV-DsRed-IRES-EGFP-GNMT 51-70 | MSCV | GNMT 51-70 | 1975 |
| CMV-DsRed-IRES-EGFP-GNMT 31-50 | MSCV | GNMT 31-50 | 1974 |
| CMV-DsRed-IRES-EGFP-GNMT 21-40 | MSCV | GNMT 21-40 | 1973 |
| CMV-DsRed-IRES-EGFP-GNMT 41-60 L50P | MSCV | GNMT 41-60 L50P | 1972 |
| CMV-DsRed-IRES-EGFP-GNMT 41-60 | MSCV | GNMT 41-60 | 1971 |
| CMV-DsRed-IRES-EGFP-GNMT 11-30 | MSCV | GNMT 11-30 | 1970 |
| CMV-DsRed-IRES-EGFP-GNMT 1-20 | MSCV | GNMT 1-20 | 1969 |
| CMV-DsRed-IRES-EGFP-GNMT 31-50 M20 | MSCV | GNMT 31-50 (Mutant Tile) | 2020 |
| CMV-DsRed-IRES-EGFP-GNMT 31-50 M19 | MSCV | GNMT 31-50 (Mutant Tile) | 2019 |
| CMV-DsRed-IRES-EGFP-GNMT 31-50 M18 | MSCV | GNMT 31-50 (Mutant Tile) | 2018 |
| CMV-DsRed-IRES-EGFP-GNMT 31-50 M17 | MSCV | GNMT 31-50 (Mutant Tile) | 2017 |
| CMV-DsRed-IRES-EGFP-GNMT 31-50 M16 | MSCV | GNMT 31-50 (Mutant Tile) | 2016 |
| CMV-DsRed-IRES-EGFP-GNMT 31-50 M15 | MSCV | GNMT 31-50 (Mutant Tile) | 2015 |
| CMV-DsRed-IRES-EGFP-GNMT 31-50 M14 | MSCV | GNMT 31-50 (Mutant Tile) | 2014 |
| CMV-DsRed-IRES-EGFP-GNMT 31-50 M13 | MSCV | GNMT 31-50 (Mutant Tile) | 2013 |
| CMV-DsRed-IRES-EGFP-GNMT 31-50 M12 | MSCV | GNMT 31-50 (Mutant Tile) | 2012 |
| CMV-DsRed-IRES-EGFP-GNMT 31-50 M11 | MSCV | GNMT 31-50 (Mutant Tile) | 2011 |
| CMV-DsRed-IRES-EGFP-GNMT 31-50 M10 | MSCV | GNMT 31-50 (Mutant Tile) | 2010 |
| CMV-DsRed-IRES-EGFP-GNMT 31-50 M9 | MSCV | GNMT 31-50 (Mutant Tile) | 2009 |
| CMV-DsRed-IRES-EGFP-GNMT 31-50 M8 | MSCV | GNMT 31-50 (Mutant Tile) | 2008 |
| CMV-DsRed-IRES-EGFP-GNMT 31-50 M7 | MSCV | GNMT 31-50 (Mutant Tile) | 2007 |
| CMV-DsRed-IRES-EGFP-GNMT 31-50 M6 | MSCV | GNMT 31-50 (Mutant Tile) | 2006 |
| CMV-DsRed-IRES-EGFP-GNMT 31-50 M5 | MSCV | GNMT 31-50 (Mutant Tile) | 2005 |
| CMV-DsRed-IRES-EGFP-GNMT 31-50 M4 | MSCV | GNMT 31-50 (Mutant Tile) | 2004 |
| CMV-DsRed-IRES-EGFP-GNMT 31-50 M3 | MSCV | GNMT 31-50 (Mutant Tile) | 2003 |
| CMV-DsRed-IRES-EGFP-GNMT 31-50 M2 | MSCV | GNMT 31-50 (Mutant Tile) | 2002 |
| CMV-DsRed-IRES-EGFP-GNMT 31-50 M1 | MSCV | GNMT 31-50 (Mutant Tile) | 2001 |
| CMV-DsRed-IRES-EGFP-GNMT 41-60 M3 | MSCV | GNMT 41-60 (Mutant Tile) | 2077 |
| CMV-DsRed-IRES-EGFP-GNMT 41-60 M15 | MSCV | GNMT 41-60 (Mutant Tile) | 2061 |
| CMV-DsRed-IRES-EGFP-GNMT 41-60 M14 | MSCV | GNMT 41-60 (Mutant Tile) | 2060 |
| CMV-DsRed-IRES-EGFP-GNMT 41-60 M13 | MSCV | GNMT 41-60 (Mutant Tile) | 2059 |
| CMV-DsRed-IRES-EGFP-GNMT 41-60 M12 | MSCV | GNMT 41-60 (Mutant Tile) | 2058 |
| CMV-DsRed-IRES-EGFP-GNMT 41-60 M11 | MSCV | GNMT 41-60 (Mutant Tile) | 2057 |
| CMV-DsRed-IRES-EGFP-GNMT 41-60 M10 | MSCV | GNMT 41-60 (Mutant Tile) | 2056 |
| CMV-DsRed-IRES-EGFP-GNMT 41-60 M9 | MSCV | GNMT 41-60 (Mutant Tile) | 2055 |
| CMV-DsRed-IRES-EGFP-GNMT 41-60 M8 | MSCV | GNMT 41-60 (Mutant Tile) | 2054 |
| CMV-DsRed-IRES-EGFP-GNMT 41-60 M7 | MSCV | GNMT 41-60 (Mutant Tile) | 2053 |
| GNMT 281-295 + C-term | GPS6.0 | GNMT 281-295 + C-term | 2063 |
| GPS6.0 CL1 | GPS6.0 | CL1 | 2051 |
| GPS6.0 GNMT 281-296 | GPS6.0 | GNMT 281-296 | 2050 |
| GPS6.0 GNMT 271-290 | GPS6.0 | GNMT 271-290 | 2049 |
| GPS6.0 GNMT 241-260 | GPS6.0 | GNMT 241-260 | 2048 |
| GPS6.0 GNMT 131-150 N141S | GPS6.0 | GNMT 131-150 N141S | 2047 |
| GPS6.0 GNMT 131-150 | GPS6.0 | GNMT 131-150 | 2046 |
| GPS6.0 GNMT 91-110 | GPS6.0 | GNMT 91-110 | 2045 |
| GPS6.0 GNMT 41-60 | GPS6.0 | GNMT 41-60 | 2044 |
| GPS6.0 GNMT 31-50 | GPS6.0 | GNMT 31-50 | 2043 |
| GPS6.0 GNMT 251-270 | GPS6.0 | GNMT 251-270 | 2076 |
| GPS6.0 GNMT 201-220 | GPS6.0 | GNMT 201-220 | 2075 |
| GPS6.0 GNMT 161-180 | GPS6.0 | GNMT 161-180 | 2074 |
| GPS6.0 GNMT 51-70 | GPS6.0 | GNMT 51-70 | 2073 |
| CMV-DsRed-IRES-EGFP-GNMT F287A | MSCV | GNMT F287A | 2325 |
| CMV-DsRed-IRES-EGFP-N141S-GNMT 271-290 single A | MSCV | N141S-GNMT 271-290 single A | 2308 |
| CMV-DsRed-IRES-EGFP-L50P-GNMT 271-290 single A | MSCV | L50P-GNMT 271-290 single A | 2307 |
| CMV-DsRed-IRES-EGFP-GNMT 271-290 single A | MSCV | GNMT 271-290 single A | 2306 |
| CMV-DsRed-IRES-EGFP-H177N-GNMT 271-290 single A | MSCV | H177N-GNMT 271-290 single A | 2298 |
| CMV-DsRed-IRES-EGFP-H177N-GNMT 41-60 single A | MSCV | H177N-GNMT 41-60 single A | 2295 |
| CMV-DsRed-IRES-EGFP-GNMT 31-50AAA + 241-260 4A + 271-290AAA | MSCV | GNMT 31-50AAA + 241-260 4A + 271-290AAA | 2290 |
| CMV-DsRed-IRES-EGFP-N141S-GNMT 41-60 single A | MSCV | N141S-GNMT 41-60 single A | 2288 |
| CMV-DsRed-IRES-EGFP-GNMT 41-60 single A | MSCV | GNMT 41-60 single A | 2287 |
| CMV-DsRed-IRES-EGFP-GNMT 31-50AAA + 271-290AAA | MSCV | GNMT 31-50AAA + 271-290AAA | 2286 |
| CMV-DsRed-IRES-EGFP-N141S-GNMT 241-260 single A | MSCV | N141S-GNMT 241-260 single A | 2285 |
| CMV-DsRed-IRES-EGFP-H177N-GNMT 241-260 single A | MSCV | H177N-GNMT 241-260 single A | 2284 |
| CMV-DsRed-IRES-EGFP-L50P-GNMT 241-260 single A | MSCV | L50P-GNMT 241-260 single A | 2283 |
| CMV-DsRed-IRES-EGFP-GNMT 241-260 single A | MSCV | GNMT 241-260 single A | 2282 |
| CMV-DsRed-IRES-EGFP-H177N-GNMT 31-50 single A | MSCV | H177N-GNMT 31-50 single A | 2281 |
| CMV-DsRed-IRES-EGFP-N141S-GNMT 31-50 single A | MSCV | N141S-GNMT 31-50 single A | 2280 |
| CMV-DsRed-IRES-EGFP-L50P-GNMT 31-50 single A | MSCV | L50P-GNMT 31-50 single A | 2279 |
| CMV-DsRed-IRES-EGFP-GNMT 31-50 single A | MSCV | GNMT 31-50 single A | 2278 |
| CMV-DsRed-IRES-EGFP-N141S-GNMT 31-50AAA + 41-60AAA + 241-260 4A + 271-290AAA | MSCV | N141S-GNMT 31-50AAA + 41-60AAA + 241-260 4A + 271-290AAA | 2277 |
| CMV-DsRed-IRES-EGFP-H177N-GNMT 271-290AAA | MSCV | H177N-GNMT 271-290AAA | 2275 |
| CMV-DsRed-IRES-EGFP-N141S-GNMT 271-290AAA | MSCV | N141S-GNMT 271-290AAA | 2274 |
| CMV-DsRed-IRES-EGFP-L50P-GNMT 271-290AAA | MSCV | L50P-GNMT 271-290AAA | 2273 |
| CMV-DsRed-IRES-EGFP-L50P-GNMT 31-50AAA + 241-260 4A + 271-290AAA | MSCV | L50P-GNMT 31-50AAA + 241-260 4A + 271-290AAA | 2248 |
| CMV-DsRed-IRES-EGFP-N141S-GNMT 31-50AAA + 41-60AAA + 241-260 4A + 271-290AAA | MSCV | N141S-GNMT 31-50AAA + 41-60AAA + 241-260 4A + 271-290AAA | 2246 |
| CMV-DsRed-IRES-EGFP-H177N-GNMT 31-50AAA + 41-60AAA + 241-260 4A + 271-290AAA | MSCV | H177N-GNMT 31-50AAA + 41-60AAA + 241-260 4A + 271-290AAA | 2245 |
| CMV-DsRed-IRES-EGFP-H177N-GNMT 31-50AAA + 41-60AAA + 271-290AAA | MSCV | H177N-GNMT 31-50AAA + 41-60AAA + 271-290AAA | 2244 |
| CMV-DsRed-IRES-EGFP-N141S-GNMT 31-50AAA + 41-60AAA + 271-290AAA | MSCV | N141S-GNMT 31-50AAA + 41-60AAA + 271-290AAA | 2243 |
| CMV-DsRed-IRES-EGFP-L50P-GNMT 31-50AAA + 271-290AAA | MSCV | L50P-GNMT 31-50AAA + 271-290AAA | 2242 |
| CMV-DsRed-IRES-EGFP-GNMT 31-50AAA + 41-60AAA + 271-290AAA | MSCV | GNMT 31-50AAA + 41-60AAA + 271-290AAA | 2241 |
| CMV-DsRed-IRES-EGFP-H177N-GNMT Full 261-260 4A | MSCV | H177N-GNMT Full 261-260 4A | 2240 |
| CMV-DsRed-IRES-EGFP-N141S-GNMT Full 261-260 4A | MSCV | N141S-GNMT Full 261-260 4A | 2239 |
| CMV-DsRed-IRES-EGFP-L50P-GNMT Full 261-260 4A | MSCV | L50P-GNMT Full 261-260 4A | 2238 |
| CMV-DsRed-IRES-EGFP-GNMT Full 261-260 4A | MSCV | GNMT Full 261-260 4A | 2237 |
| CMV-DsRed-IRES-EGFP-H177N-GNMT 31-50AAA + 41-60AAA | MSCV | H177N-GNMT 31-50AAA + 41-60AAA | 2233 |
| CMV-DsRed-IRES-EGFP-N141S-GNMT 31-50AAA + 41-60AAA | MSCV | N141S-GNMT 31-50AAA + 41-60AAA | 2232 |
| CMV-DsRed-IRES-EGFP-GNMT 31-50AAA + 41-60AAA | MSCV | GNMT 31-50AAA + 41-60AAA | 2231 |
| CMV-DsRed-IRES-EGFP-H177N-GNMT Full 31-50AAA | MSCV | H177N-GNMT Full 31-50AAA | 2227 |
| CMV-DsRed-IRES-EGFP-H177N-GNMT Full 41-60AAA | MSCV | H177N-GNMT Full 41-60AAA | 2226 |
| CMV-DsRed-IRES-EGFP-N140S-GNMT Full 41-60AAA | MSCV | N140S-GNMT Full 41-60AAA | 2225 |
| CMV-DsRed-IRES-EGFP-N140S-GNMT Full 31-50AAA | MSCV | N140S-GNMT Full 31-50AAA | 2224 |
| CMV-DsRed-IRES-EGFP-L50P-GNMT Full 31-50AAA | MSCV | L50P-GNMT Full 31-50AAA | 2223 |

**Supplementary Table S4: siRNA sequences and catalog numbers**

| **Gene Symbol** | **Full Gene Name** | **Manufacturer** | **Catalog Number** | **Sense Sequence** | **Antisense Sequence** | **Target Sequence for SMARTpool** |
| --- | --- | --- | --- | --- | --- | --- |
| AIP | aryl hydrocarbon receptor interacting protein | Thermo Fisher Scientific (Silencer Select Series) | s531246 | GGGCCAUGACAGACGAAGAtt | UCUUCGUCUGUCAUGGCCCat | NA |
| ALG13 | asparagine-linked glycosylation 13 homolog | Thermo Fisher Scientific (Silencer Select Series) | s31656 | CAAGGAUUCCUUGAAAGAAtt | UUCUUUCAAGGAAUCCUUGta | NA |
| BAG2 | BCL2-associated athanogene 2 | Thermo Fisher Scientific (Silencer Select Series) | s18293 | GCAAGAAUCCCUAAAGCAUtt | AUGCUUUAGGGAUUCUUGCtg | NA |
| BAT3 | HLA-B associated transcript 3 | Thermo Fisher Scientific (Silencer Select Series) | s15466 | CCUUCAAUCUUCCUAGUGAtt | UCACUAGGAAGAUUGAAGGtt | NA |
| BAT3 | HLA-B associated transcript 3 | Thermo Fisher Scientific (Silencer Select Series) | s15468 | CAAGAGCAGUUUAAUAGCAtt | UGCUAUUAAACUGCUCUUGga | NA |
| BAT3 | HLA-B associated transcript 3 | Dharmacon (ON-TARGETplus siRNA - SMARTpool) | L-005062-01-0005 | Sequences not provided | Sequences not provided | GAGGAGGAUCAGCGGUUGA; UGUUAUCAAUGGCCGAAUU; UCUCUAUGGUGGACGUAGU; ACAUUCAGAGCCAGCGGAA |
| BIRC6 | baculoviral IAP repeat-containing 6 | Thermo Fisher Scientific (Silencer Select Series) | s33037 | GGUAGAUAUUUCUAGUACAtt | UGUACUAGAAAUAUCUACCtt | NA |
| CACYBP | calcyclin binding protein | Thermo Fisher Scientific (Silencer Select Series) | s25819 | CCAUGAUUGUGAACAAUCUtt | AGAUUGUUCACAAUCAUGGag | NA |
| DMWD | dystrophia myotonica, WD repeat containing | Thermo Fisher Scientific (Silencer Select Series) | s4171 | GUACCUGGAUCUCAUCAAAtt | UUUGAUGAGAUCCAGGUACtg | NA |
| DNAJA1 | DnaJ (Hsp40) homolog, subfamily A, member 1 | Thermo Fisher Scientific (Silencer Select Series) | s6964 | GGAAUUAUAUGACAAAGGAtt | UCCUUUGUCAUAUAAUUCCct | NA |
| DNAJA2 | DnaJ (Hsp40) homolog, subfamily A, member 2 | Thermo Fisher Scientific (Silencer Select Series) | s20124 | CAAUCAGAGUAGAAGUCGAtt | UCGACUUCUACUCUGAUUGcc | NA |
| DNAJC7 | DnaJ (Hsp40) homolog, subfamily C, member 7 | Thermo Fisher Scientific (Silencer Select Series) | s14473 | CCCUAGAACUGGAUCAUAAtt | UUAUGAUCCAGUUCUAGGGct | NA |
| DZIP3 | DAZ interacting protein 3, zinc finger | Thermo Fisher Scientific (Silencer Select Series) | s18597 | CAGGUGACAUGGUAAGGAUtt | AUCCUUACCAUGUCACCUGta | NA |
| FBOX21 | F-box protein 21 | Dharmacon (ON-TARGETplus siRNA - SMARTpool) | L-012917-00-0005 | Sequences not provided | Sequences not provided | GAACAUAGAUGAGUAAAGU; UCAAUUGGUUGGAAGAGUA; AGGAUGAACUGGUGUGUAU; UCUUGAAGGUGCUGUAUAU |
| FEM1B | Fem-1 homolog B | Dharmacon (ON-TARGETplus siRNA - SMARTpool) | L-015838-00-0005 | Sequences not provided | Sequences not provided | ACACUGACAUGACGAAUAA; GCCUAAUGAUUGCGGCAUA; AGAAUAAGACUCCGCUAGA; ACAAUGCUAUGGACAAUUA |
| FKBP5 | FK506 binding protein 5 | Thermo Fisher Scientific (Silencer Select Series) | s5215 | GAGAAAGGCUUGUAUAGGAtt | UCCUAUACAAGCCUUUCUCat | NA |
| GET4 | golgi to ER traffic protein 4 homolog | Thermo Fisher Scientific (Silencer Select Series) | s28405 | AGAUGUACCGGACCCUGUUtt | AACAGGGUCCGGUACAUCUgg | NA |
| GET4 | golgi to ER traffic protein 4 homolog | Thermo Fisher Scientific (Silencer Select Series) | s28406 | CGAGUACCUCGACCGCAUAtt | UAUGCGGUCGAGGUACUCGtt | NA |
| KLHL8 | kelch-like family member 8 | Thermo Fisher Scientific (Silencer Select Series) | s33321 | CGAAUAGACUUAAUGGACAtt | UGUCCAUUAAGUCUAUUCGat | NA |
| NA | ON-TARGETplus Non-targeting Pool | Dharmacon (ON-TARGETplus siRNA - SMARTpool) | D-001810-10-05 | Sequences not provided | Sequences not provided | NA |
| OTUD4 | OTU domain containing 4 | Thermo Fisher Scientific (Silencer Select Series) | s29347 | UAAUUUAUCGGGAACCAAAtt | UUUGGUUCCCGAUAAAUUAta | NA |
| PDZRN3 | PDZ domain containing ring finger 3 | Thermo Fisher Scientific (Silencer Select Series) | s22854 | GGUCAACGGCAGAGACUUAtt | UAAGUCUCUGCCGUUGACCtc | NA |
| PJA2 | praja ring finger 2 | Thermo Fisher Scientific (Silencer Select Series) | s19106 | CAGGAAUCAUUAGGCAAUAtt | UAUUGCCUAAUGAUUCCUGgt | NA |
| PTGES3 | prostaglandin E synthase 3 (cytosolic) | Thermo Fisher Scientific (Silencer Select Series) | s21075 | CGACUUCAAUAAUUGGAAAtt | UUUCCAAUUAUUGAAGUCGac | NA |
| RNF126 | ring finger protein 126 | Dharmacon (ON-TARGETplus siRNA - SMARTpool) | L-007015-00-0005 | Sequences not provided | Sequences not provided | UGUCUAACCUCACCCUCUA; CAUCACACAGCUCCUCAAU; CGGAUUAUAUCUGUCCAAG; GAACAAAACUGCUCCAACA |
| SGTA | small glutamine-rich tetratricopeptide repeat (TPR)-containing, alpha | Thermo Fisher Scientific (Silencer Select Series) | s12781 | GAAGGAAACGAGCAGAUGAtt | UCAUCUGCUCGUUUCCUUCgg | NA |
| SGTA | small glutamine-rich tetratricopeptide repeat (TPR)-containing, alpha | Thermo Fisher Scientific (Silencer Select Series) | s12783 | GCUUCGAACCUAAUGAACAtt | UGUUCAUUAGGUUCGAAGCca | NA |
| Negative Control - #1 | NA | Thermo Fisher Scientific (Silencer Select Series) | s813 | Sequence Not provided | Sequence Not provided | NA |
| Negative Control - #2 | NA | Thermo Fisher Scientific (Silencer Select Series) | s814 | Sequence Not provided | Sequence Not provided | NA |
| SKP1 | S-phase kinase-associated protein 1 | Thermo Fisher Scientific (Silencer Select Series) | s12890 | AACAAUCUGUGACUAUUAAtt | UUAAUAGUCACAGAUUGUUtg | NA |
| SQSTM1 | sequestosome 1 | Thermo Fisher Scientific (Silencer Select Series) | s16960 | GGAGCACGGAGGGAAAAGAtt | UCUUUUCCCUCCGUGCUCCac | NA |
| ST13 | suppression of tumorigenicity 13 (colon carcinoma) (Hsp70 interacting protein) | Thermo Fisher Scientific (Silencer Select Series) | s13516 | GAGCAUGGGUGGUAAAGUAtt | UACUUUACCACCCAUGCUCtc | NA |
| STUB1 | STIP1 homology and U-box containing protein 1, E3 ubiquitin protein ligase | Thermo Fisher Scientific (Silencer Select Series) | s195026 | CGCUGGUGGCCGUGUAUUAtt | UAAUACACGGCCACCAGCGgg | NA |
| SUGT1 | SGT1, suppressor of G2 allele of SKP1 | Thermo Fisher Scientific (Silencer Select Series) | s21446 | GGUGUGGACUCAUCAGUCAtt | UGACUGAUGAGUCCACACCtc | NA |
| TRIM37 | tripartite motif containing 37 | Thermo Fisher Scientific (Silencer Select Series) | s534001 | GGGUUCCAGUAGUAGAAUAtt | UAUUCUACUACUGGAACCCac | NA |
| TRIM62 | tripartite motif-containing 62 | Thermo Fisher Scientific (Silencer Select Series) | s30482 | GGGACAAGCUUGACAAGGUtt | ACCUUGUCAAGCUUGUCCCgg | NA |
| TTC1 | tetratricopeptide repeat domain 1 | Thermo Fisher Scientific (Silencer Select Series) | s14469 | GGAAGACUAUAAAUCUAUAtt | UAUAGAUUUAUAGUCUUCCag | NA |
| UBL4A | ubiquitin-like 4A | Thermo Fisher Scientific (Silencer Select Series) | s15760 | CAACUCCAAGCUCAACCUAtt | UAGGUUGAGCUUGGAGUUGgg | NA |
| UBL4A | ubiquitin-like 4A | Thermo Fisher Scientific (Silencer Select Series) | s15761 | GCAUGUCAUGUUUAGCCCAtt | UGGGCUAAACAUGACAUGCga | NA |
| USP19 | ubiquitin specific peptidase 19 | Thermo Fisher Scientific (Silencer Select Series) | s21341 | CCAUCACUUUUGACCCGUUtt | AACGGGUCAAAAGUGAUGGag | NA |

**Supplementary Table 5: Antibodies used in this study**

| **Antibody** | **Species raised in** | **Manufacturer** | **Catalog Number** | **Western Blot/Immunofluorescence** | **Dilution** |
| --- | --- | --- | --- | --- | --- |
| BAG6 | Mouse | Santa Cruz | SC-365928 | WB | 1 in 500 |
| RNF126 | Mouse | Proteintech | 66647-1-Ig | WB | 1 in 1000 |
| Streptavidin-IR Dye 800CW | NA | Licor | 926-32230 | WB | 1 in 10,000 |
| V5 | Mouse | Invitrogen | R960-25 | Both | 1 in 1000 for WB; 1in 100 for IF |
| Biotin | Rabbit | Abcam | AB53494 | IF | 1 in 100 |
| GAPDH | Mouse | AbLab | AB000BE | WB | 1 in 5000 |
| Alpha-Tubulin | Mouse | Proteintech | 66031-1 | WB | 1 in 1000 |
| FBXO21 | Mouse | OriGENE | TA504013S | WB | 1 in 1000 |
| FEM1B | Rabbit | Proteintech | 19544-1-AP | WB | 1 in 1000 |
| Alexa Fluor-488 anti-rabbit | Goat | Invitrogen | A11008 | IF | 1 in 400 |
| Alexa Fluor-594 anti-mouse | Goat | Invitrogen | A11020 | IF | 1 in 400 |
| IRDye 800CW anti-rabbit IgG | Goat | Licor | 925-32211 | WB | 1 in 10,000 |
| IRDye 680RD anti-mouse IgG | Goat | Licor | 925-68070 | WB | 1 in 10,000 |
| IRDye 680RD anti-rabbit IgG | Goat | Licor | 926-68071 | WB | 1 in 10,000 |
| IRDye 800CW anti-mouse IgG | Goat | Licor | 926-32210 | WB | 1 in 10,000 |

**Supplementary Table 6:** **Description of features taken as input for support vector regression models.**

| **Feature** | **Description** |
| --- | --- |
| Amino Acid Fractional Composition | $\frac{Count of specific residue}{Sequence length}$ for each amino acid |
| Fraction Squared Sum | $\sum_{i = 1}^{20} x_{i}^{2}$  where x_i_ is the amino acid fractional composition for each amino acid *i* |
| Repeat Score | $\sum_{k = 2}^{L} k\times(\# of unique k-mers)$  where L is the length of the input sequence and only k-mers of disorder promoting amino acids are considered (A, R, G, Q, E, K, P, S) |
| BAG6 Motif | ≥4 of (L, I, V, M) within a 6 residue window |
| Amphipathic | ΦxxΦxxxΦ or ΦxxxΦxxΦ, where Φ = (L, I, V, F, Y, W) |
| Largest Motif Score | $\sum_{i = 0}^{L} {-\rho}_{i}$  where *L* is the length of continuous sequence that maximizes the score and *ρ*_i_ is the Pearson correlation coefficient between amino acid fractional composition and protein stability index |
| Largest Motif Length | Value of *L* used for Largest Motif Score |
| Flanking Residues | Includes flanking residues in calculating above features |

**Supplementary Table 7. AlphaFold calculations**

|  |  | **GNMT Active** | | **GNMT ΔAAA** | |  |
| --- | --- | --- | --- | --- | --- | --- |
| **Bag6 domain(s)** | **residue numbering** | **ipTM** | **pTM** | **ipTM** | **pTM** | **∆ ipTM** |
| UBL | 16-88 | 0.8 | 0.89 | 0.78 | 0.88 | 0.02 |
| Domain I | 1-240 | 0.45 | 0.38 | 0.5 | 0.37 | -0.05 |
| **BUILD** | **93-220** | **0.74** | **0.47** | **0.42** | **0.35** | **0.32** |
| Shorten Domain I | 1-195 | 0.43 | 0.41 | 0.38 | 0.41 | 0.05 |
| **IDR1 and Coil of BUILD** | **88-158** | **0.66** | **0.4** | **0.2** | **0.16** | **0.46** |
| BUILD and Domain II | 88-471 | 0.5 | 0.43 | 0.55 | 0.43 | -0.05 |
| Helix, BUILD, IDR2 and Domain II | 160-364 | 0.54 | 0.59 | 0.63 | 0.59 | -0.09 |
| Domain II | 220-471 | 0.41 | 0.41 | 0.4 | 0.41 | 0.01 |
| IDR2 in Domain II | 191-272 | 0.29 | 0.14 | 0.3 | 0.16 | -0.01 |
| Domain II and IDR3 | 386-529 | 0.29 | 0.17 | 0.33 | 0.17 | -0.04 |
| Proline Rich II Domain | 533-937 | 0.42 | 0.55 | 0.67 | 0.53 | -0.25 |
| Shorter Pro-rich domain (no IDR) | 697-937 | 0.37 | 0.68 | 0.65 | 0.76 | -0.28 |
| IDR4 | 562-690 | 0.33 | 0.24 | 0.35 | 0.25 | -0.02 |
| IDR5 | 950-1008 | 0.45 | 0.43 | 0.46 | 0.42 | -0.01 |
| BAG | 1048-1132 | 0.53 | 0.45 | 0.52 | 0.48 | 0.01 |
| C-term NLS BAG | 1008-1132 | 0.38 | 0.31 | 0.32 | 0.29 | 0.06 |
